# Effect of tRNA maturase depletion on the levels and stabilities of ribosome assembly cofactor mRNAs in *Bacillus subtilis*

**DOI:** 10.1101/2022.11.23.517781

**Authors:** Aude Trinquier, Ciarán Condon, Frédérique Braun

**Affiliations:** CNRS, Université Paris Cité, Expression Génétique Microbienne, Institut de Biologie Physico-Chimique, 13 rue Pierre et Marie Curie, 75005 Paris, France

**Keywords:** RNA degradation, tRNA processing, translation, chloramphenicol, (p)ppGpp

## Abstract

The impact of translation on mRNA stability can be varied, ranging from a protective effect of ribosomes that shield mRNA from ribonucleases (RNases), to preferentially exposing sites of RNase cleavage. These effects can change depending on whether ribosomes are actively moving along the mRNA or whether they are stalled at particular sequences, structures or awaiting charged tRNAs. We recently observed that depleting *B. subtilis* cells of its tRNA maturation enzymes RNase P or RNase Z, led to altered mRNA levels of a number of assembly factors involved in the biogenesis of the 30S ribosomal subunit. Here, we extend this study to other assembly factor mRNAs and identify multiple transcriptional and translational layers of regulation of the *rimM* operon mRNA that occur in response to the depletion of functional tRNAs.

**Importance:** The passage of ribosomes across individual mRNAs during translation can have different effects on their degradation, ranging from a protective effect by shielding from ribonucleases, to in some cases, making the mRNA more vulnerable to RNase action. We recently showed that some mRNAs coding for proteins involved in ribosome assembly were highly sensitive to the availability of functional tRNA. Using strains depleted for the major tRNA processing enzymes RNase P and RNase Z, we expanded this observation to a wider set of mRNAs, including some unrelated to ribosome biogenesis. We characterize the impact of tRNA maturase depletion on the *rimM* operon mRNA and show it is highly complex, with multiple levels of transcriptional and post-transcriptional effects coming into play.

## Introduction

The steady-state level of any mRNA in the cell is determined by both its rate of transcription by RNA polymerase and its rate of degradation by ribonucleases (RNases). These can work together to increase or decrease gene expression at the transcriptional or post-transcriptional levels in response to environmental stimuli, or can pull in opposing directions, resulting in little net gain. While most of the enzymes responsible for RNA decay are now known in *B. subtilis* (1), how these enzymes are impacted by translation is still a relatively open question. The conventional wisdom is that increased translation leads to increased stability due to the masking of RNase cleavage sites by ribosomes. However, we have recently identified an endoribonuclease (Rae1) that actually depends on translation to destabilize mRNAs (2, 3), and we will present data here suggesting that antibiotics that cause ribosome pausing can both positively and negatively impact mRNA levels, depending on the severity and time of the dose.

Efficient translation depends on an unlimited supply of functional charged transfer RNAs (tRNAs). tRNAs are almost universally transcribed as precursors in all living organisms, with both 5’ and 3’ extensions that must be removed to generate tRNAs that can be charged and used in translation. In *B. subtilis*, tRNAs are matured at their 5’ end by the ubiquitous endoribonuclease RNase P, consisting of a catalytic RNA moiety encoded by the *rnpB* gene and a protein subunit encoded by the *rnpA* gene (4). Their 3’ ends are matured either by the endo/exoribonuclease RNase Z, or by a number of redundant 3’ exoribonucleases, depending nominally on whether the tRNA gene encodes the CCA motif required for aminoacylation (5). RNase Z processes the one-third of *B. subtilis* tRNAs lacking an encoded CCA-motif through stimulation of its endoribonuclease activity about 200-fold by a uracil residue that naturally occurs ≤ 2 nts downstream of the so-called discriminator nucleotide (nt 75) of each of these tRNA precursors (6). The enzyme’s 3’-exoribonuclease activity is required to trim back to nt 75 to allow addition of the CCA motif by nucleotidyl-transferase (NTase or CCAse). Both RNase P and Z are essential in *B. subtilis* and depletion of either enzyme inhibits cell growth, presumably due to a lack of functional tRNAs for translation.

Translation can also be inhibited by antibiotics that target the ribosome, such as chloramphenicol (Cm), that that targets the peptidyl transferase center (PTC) located on the large ribosomal subunit. Although it was once thought that Cm blocked translation randomly, recent ribosome profiling experiments have shown that Cm preferentially causes ribosomes to stall at particular sites, in particular when alanine (Ala) or serine (Ser) residues have just been incorporated into the nascent peptide (7).

We have recently shown that depletion of tRNA maturase activity affects ribosome assembly leading to a specific 30S subunit late assembly defect (8). While this defect was mostly explained by a RelA-dependent accumulation of the stringent response alarmone (p)ppGpp, and inhibition of GTP-dependent assembly factor activity, we also observed that the levels of several mRNAs encoding ribosome assembly cofactors were affected. Notably, the steady state levels of transcripts encoding the GTPases Era and YqeH were up-regulated during tRNA maturase depletion, whereas mRNAs encoding the GTPase CpgA and the RNA chaperone RimM were down-regulated. Because RNase P is thought to have very few direct mRNA targets, and because RNase Z depletion had comparable effects on the expression of these mRNAs, we considered it unlikely that the effects observed were directly due to RNase P or RNase Z cleavages in each of these mRNAs. We therefore wished to better understand by which mechanism(s) tRNA maturase depletion affected the levels of the cofactor encoding mRNAs. Since the late 30S ribosome assembly defect observed in tRNA maturase depletion strains was very similar to that observed in both *E. coli* and *B. subtilis ΔrimM* mutants, we put additional focus on exploring the decrease in *rimM* expression under these conditions.

## Results

### tRNA maturase depletion alters assembly factor mRNA levels

We previously showed that depletion of RNase P or RNase Z results in altered mRNA levels of four key 30S assembly cofactors (Era, YqeH, RimM and CpgA) (8). The effects of depleting the RNA subunit of RNase P (RnpB) were more severe than the protein subunit (RnpA), presumably because the RNA component of RNase P is more rapidly depleted than the protein subunit once transcription is shut off. To ask whether this applied to other mRNAs involved in ribosome biogenesis, we extended this analysis to the expression of several other cofactor and ribosomal protein mRNAs, using xylose (*Pxyl-rnpA*) or IPTG-dependent (*Pspac-rnpB* and *Pspac-rnz*) promoter constructs to deplete the protein and RNA subunits of RNase P, and RNase Z, respectively. The two control transcripts, *yqeH* and *era*, and three new transcripts, *ydaF* and *yjcK* (encoding two potential homologs of the *E. coli* RimJ acetylase), and *rpsU* (encoding r-protein S21) were globally increased under conditions of tRNA maturase depletion (Figure 1A), while *rimM, cpgA* (controls) and *yfmL* transcripts (encoding a DEAD-box helicase), all showed decreased expression, with a visible accumulation of degradation intermediates for *yfmL* (Figure 1B). Expression of the *rbfA* and *ylxS/rimP* mRNAs were relatively unchanged (Figure 1C), showing that tRNA maturase depletion does not cause non-specific perturbation of the expression of all *B. subtilis* ribosome assembly cofactor genes. Although the primary focus of this study was on assembly factor mRNAs because of the link to a defect in 30S biogenesis, we also asked whether effects of tRNA maturase depletion could be seen on other mRNAs. Indeed, mRNAs from the *yrzI* and *bmrCD* operons, encoding multiple peptides of unknown function and a multi-drug resistance pump, respectively, were also up-regulated upon RNase P or RNase Z depletion (Figure 1D), suggesting this phenomenon is not confined to mRNAs with ribosome-related functions.

**Figure 1.**
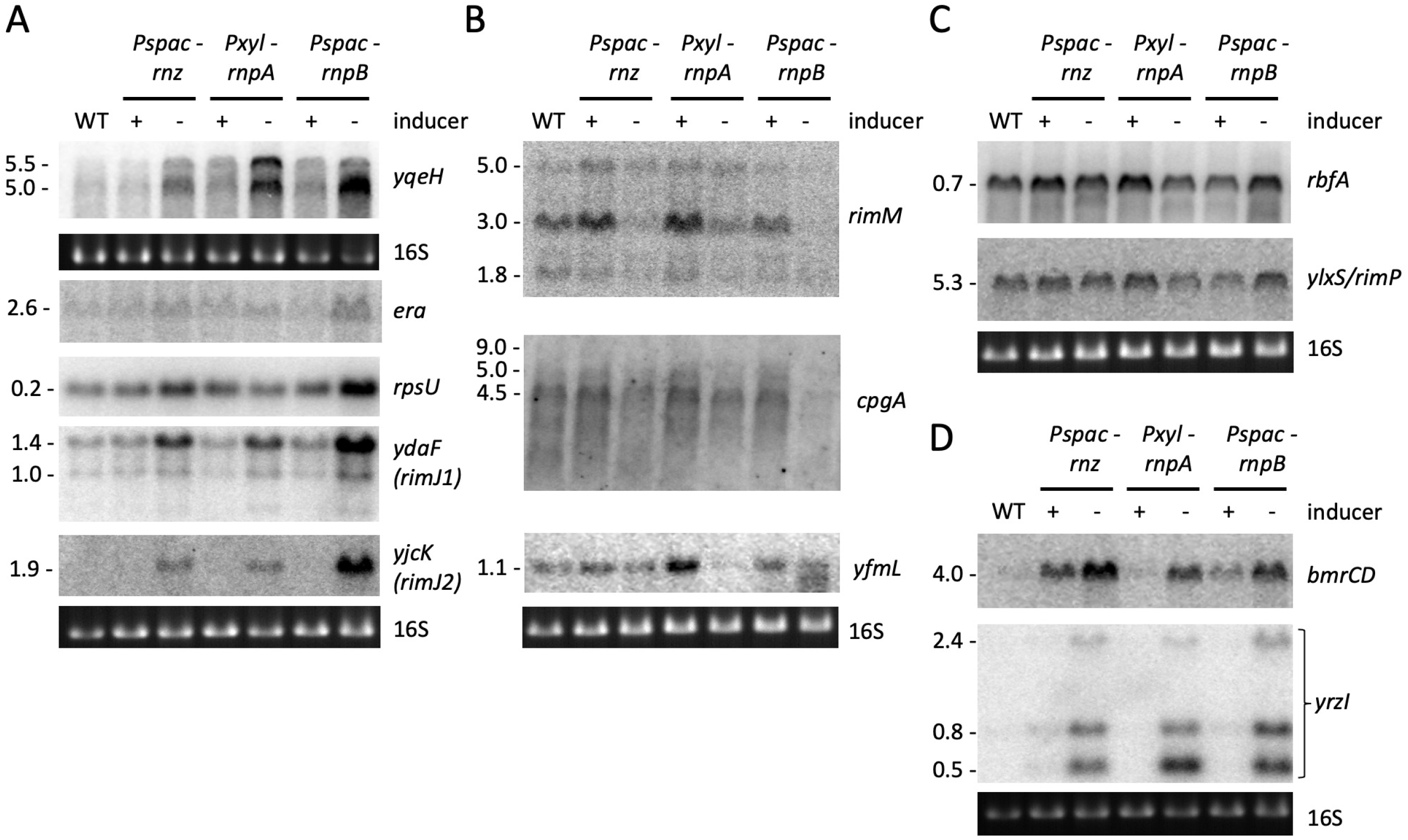
Depletion of tRNA processing enzymes results in perturbed expression of some mRNAs encoding proteins involved in 30S subunit assembly. Northern blots showing (A) Up-regulated mRNAs, (B) Down-regulated mRNAs (C) Unaffected mRNAs and (D) mRNAs unrelated to ribosome assembly, present in total mRNA isolated in the presence or absence of inducer as indicated. Note that the basal level of the *bmrCD* transcript, encoding a multidrug transporter, is higher in the *Pspac-rnz* and *Pspac-rnpB* strains because of the presence of erythromycin in the medium for stable maintenance of the construct. 16S rRNA levels (ethidium bromide stained) are shown as a loading control. Series of blots where a single loading control is shown, were stripped and reprobed. The blots for *era, yqeH, rimM* and *cpgA* were regenerated as in ref. (8) with independent RNA preparations, with permission granted by the publisher for re-use of previously published data. Number of repetitions (n) as follows: *yqeH* (n=3); *era* (n=3); *rpsU* (n=4); *ydaF* (n=2); *yjcK* (n=2); *rimM* (n=3); *cpgA* (n=3); *rbfA* (n=2); *ylxS* (n=2); *yfmL* (n=2); *bmrCD* (n=2); *yrzI* (n=2).

### tRNA maturase depletion and the translation inhibitor chloramphenicol alter mRNA stability in a similar manner

To determine whether tRNA maturase depletion impacted mRNA expression at the transcriptional or post-transcriptional level, we measured the stability of several of these mRNAs after rifampicin treatment in RNase P (RnpA or RnpB) depleted cultures. The up-regulated transcripts (*yqeH, era, ydaF* and *yjcK*) were all stabilized during RnpA and RnpB depletion (Figure 2A and B; Figure S1), suggesting that they are affected by RNase P depletion at the post-transcriptional level. There is evidence that the lack of functional tRNAs can increase ribosome stalling on translated mRNAs (9). Thus, the increased stability of these transcripts could be due to ribosome stalling and the blocking of ribonuclease access to cleavage sites on these mRNAs. To test this hypothesis, we sought to recapitulate the effect by pausing translation in a different manner, using the translation elongation inhibitor Cm. Indeed, the addition of sub-inhibitory (2.5 μg/mL) and minimal inhibitory (MIC; 5 μg/mL) concentrations of Cm to WT cells also increased the levels of the *yqeH, era, ydaF* and *yjcK* mRNAs (Figure 2C), suggesting that the stabilization of these transcripts in tRNA maturase depletion strains is most likely due to the lack of mature tRNA and ribosome stalling.

**Figure 2.**
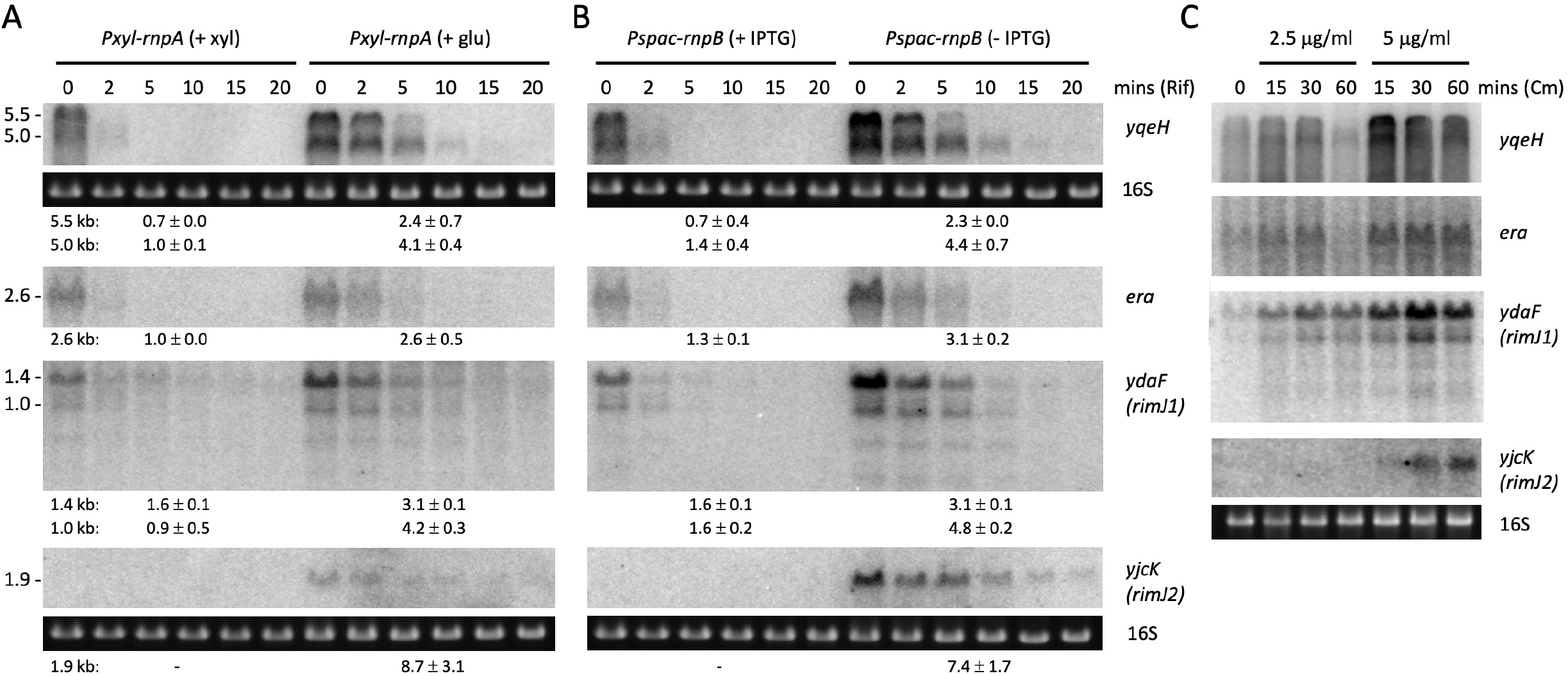
Up-regulated mRNAs show increased stability upon tRNA maturase depletion and increased expression levels in the presence of chloramphenicol. Northern blots of total RNA isolated at different times after addition of rifampicin (Rif) in cells grown in the presence or absence of inducer for (A) *rnpB* or (B) *rnpA* expression. Transcript sizes are given in kb to the left of the blots and half-lives are reported under each blot. Note, that since *yjcK* gives no signal in the presence of inducer, we cannot rule out a transcriptional effect in this case. (C) Northern blots of total RNA isolated at different times after addition of 0.5x MIC and MIC of chloramphenicol (Cm). 16S rRNA levels (ethidium bromide stained) are shown as a loading control. Series of blots where a single loading control is shown, were stripped and reprobed. Experiments were performed twice, with decay plots and their quantifications given in Figure S1.

The situation with the down-regulated transcripts was more complicated. The major *rimM* (5, 3.5 and 1.8 kb) and *cpgA* (5 and 4.5 kb) transcripts were strongly destabilized in RNase P RNA subunit (RnpB) depleted cells (Figure 3B; Figure S2), suggesting that the decrease in expression also occurs at a post-transcriptional level in this strain. A similar decrease in expression was seen after 30 minutes at high (MIC) Cm concentration in WT cells for *rimM* and rapidly upon exposure to Cm for *cpgA* (Figure 3C), suggesting that this phenomenon is also linked to ribosome stalling. For both *cpgA* and *yfmL*, the major transcripts were processed to shorter forms in the absence of RnpB or in the presence of Cm (Fig. 3B). One possibility is that, in contrast to the up-regulated mRNAs, when non-functional tRNA precursors accumulate to high levels in the *rnpB*-depletion strain, ribosomes eventually stall at sites that preferentially allow RNase access, or that the RNAs are largely unoccupied by ribosomes.

**Figure 3.**
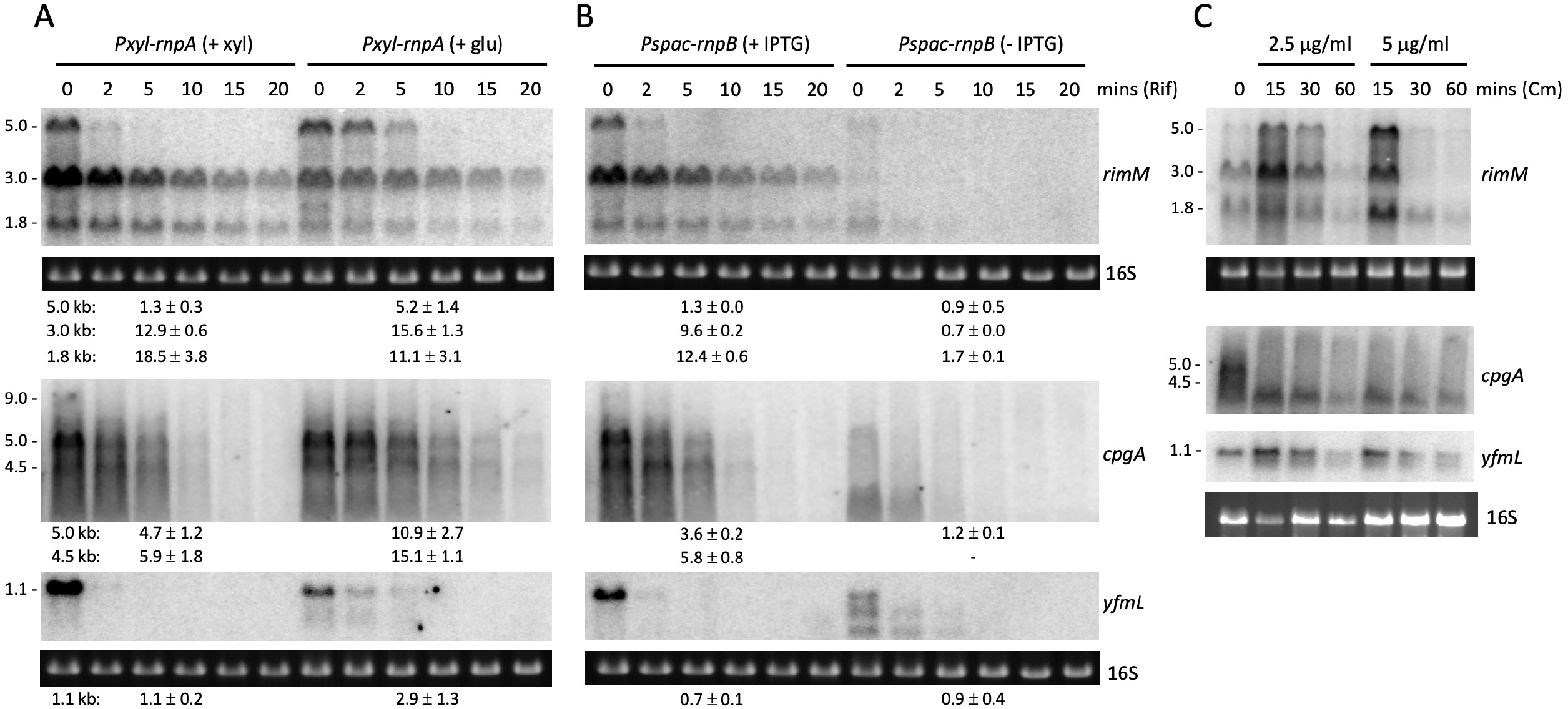
Downregulated mRNAs are subjected to a mixture of transcriptional and post-transcriptional effects upon tRNA maturase depletion and chloramphenicol addition. Northern blots of total RNA isolated at different times after addition of rifampicin (Rif) in cells grown in the presence or absence of inducer for (A) *rnpB* or (B) *rnpA* expression. Transcript sizes are given in kb to the left of the blot and half-lives are reported under each blot. (C) Northern blots of total RNA isolated at different times after addition of 0.5x MIC and MIC of chloramphenicol (Cm). 16S rRNA levels (ethidium bromide stained) are shown as a loading control. Series of blots where a single loading control is shown, were stripped and reprobed. Blots shown in Fig. 2 were stripped and reprobed for use in this figure. Experiments were performed twice, with decay plots and their quantifications given in Figure S2.

In the less severely depleted *rnpA* strain, the full-length (5 kb) *rimM* transcript and the two major *cpgA* mRNAs were stabilized (or showed little effect), rather than destabilized as seen for *rnpB* (Figure 3A). These results suggest that down-regulation of *rimM* and *cpgA* arises from a mixture of transcriptional (down) and post-transcriptional (up initially, then down) effects and that one or other effect predominates depending on the severity of RNase P depletion. Indeed, upon close inspection of Figure 3C, *rimM* and *cpgA* mRNA levels initially increase at 15 mins and then decrease after further exposure to Cm at both sub-inhibitory and MIC doses. Thus, the Cm effect globally tracks the effect of tRNA depletion, with the weak Cm dose (2.5 μg/mL) mimicking the weak effect of depleting RnpA, and the strong Cm dose (5 μg/mL) mimicking the strong effect of depleting RnpB, consistent with the notion of opposing responses to severe *vs* less severe levels or duration of translation inhibition.

### Identification of *rimM*-containing transcripts sensitive to RNase P depletion

Because the *ΔrimM* phenotype closely fitted the 30S late assembly defect observed in strains depleted for RNase P or RNase Z (8), we attempted to narrow down the determinants of the down-regulation of this operon. The *rimM* gene is encoded in a large operon containing several genes encoding components of the translation machinery: ribosomal protein genes *rpsP* and *rplS* (encoding S16 and L19, respectively), signal recognition particle components (encoded by *ffh* and *ylxM*) and *trmD* that encodes a tRNA methyltransferase. To identify the gene composition of the three *rimM-*containing transcripts, we performed northern blots with probes located in ORFs of the neighboring genes (Figure S3). In all, six different transcripts originate from this locus (Figure 4A). Promoters upstream of *ylxM* (P_1_) and *rplS* (P_3_), and terminators downstream of *ylqC* and *rplS* (T_1_ and T_3_, respectively) were identified earlier by transcriptome analysis (10). Our Northern blot analysis suggested that two transcripts originate from P_1_: the full-length mRNA (5 kb, highlighted in purple) that terminates at T_3_, and a shorter transcript (2.5 kb, highlighted in orange) that terminates at T_1_ and does not contain the *rimM* ORF. The smallest species identified (0.5 kb, highlighted in yellow) corresponds to the mono-cistronic *rplS* transcript (P_3_ to T_3_). Using end-enrichment RNA sequencing (Rend-seq), DeLoughery *et al*. identified a third transcription start site (P_2_) located just upstream of *rpsP* and only 18 nts downstream of an RNase Y cleavage site in the *ffh-rpsP* intergenic region, in addition to a potential terminator/attenuator (T_2_) within the *trmD* ORF (11). The three remaining transcripts (0.7 kb, highlighted in green; 1.8 kb, in cyan; 3 kb, in pink, and marked with an asterisk in Figure 4A) therefore correspond either to P_2_ primary transcripts, or RNase Y-processed transcripts originating from P_1_, which terminate at T_1_, T_2_ and T_3_, respectively. Interestingly, of the six transcripts encoded by this locus, only the three containing both the *ylqD* and *rimM* ORFs were down-regulated upon RNase P depletion (Figure 4B).

**Figure 4.**
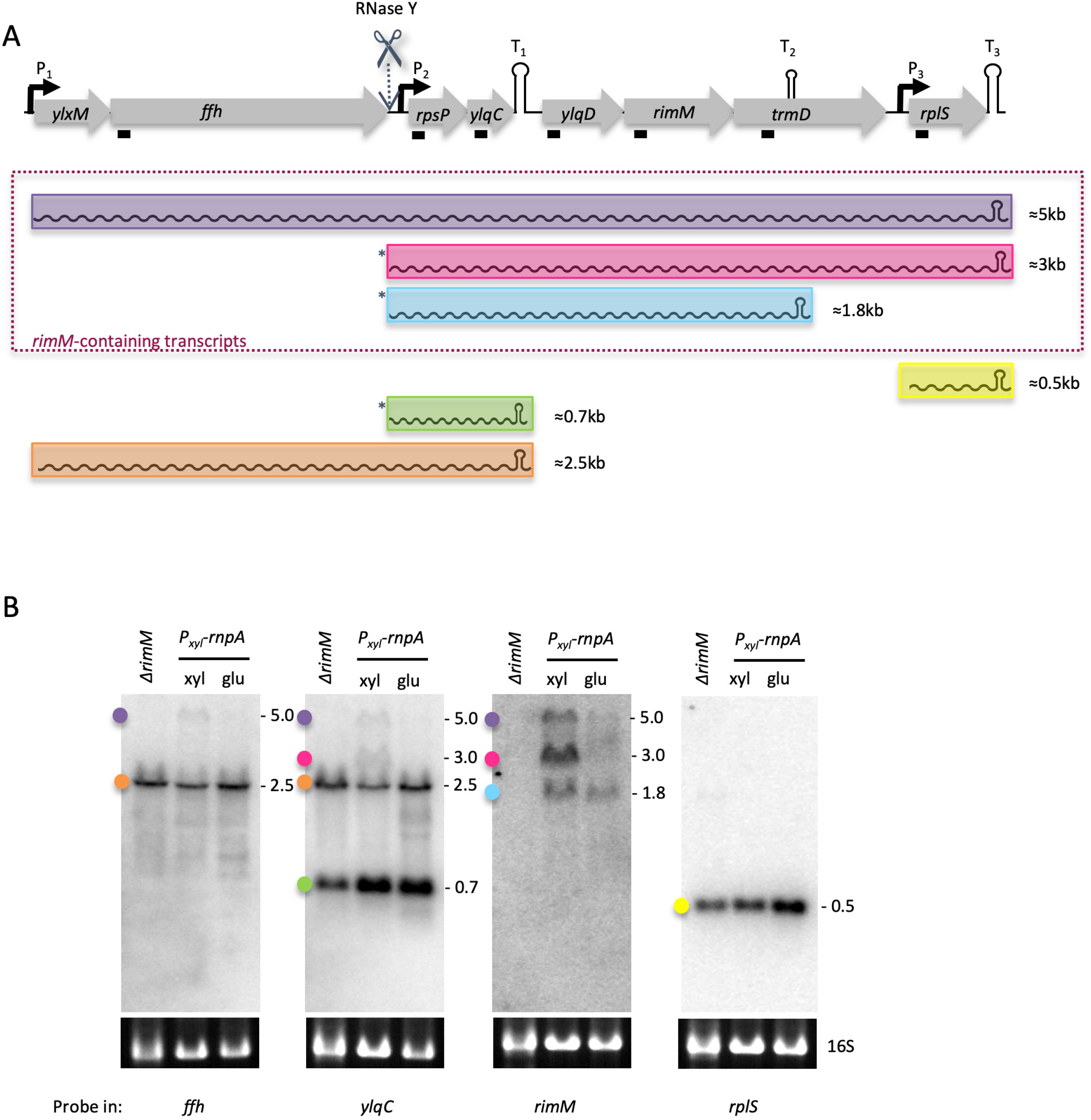
Three of the six transcripts emanating from the *rimM* operon are sensitive to tRNA maturase depletion. (A) Structure of the *rimM* operon. Open reading frames (ORFs; not to scale) are shown as gray arrows and transcripts as different colored wavy lines. Sizes are as indicated. Promoters (P_1_-P_3_) are represented by black arrows and terminators (T_1_-T_3_) as hairpins. A known RNase Y cleavage site is indicated by a scissors symbol. The asterisk indicates transcripts that may be processed by RNase Y, but are not distinguishable from P2 primary transcripts by Northern blot. These are denoted Y/P2 in the text. (B) Northern blot analysis of total RNA from *Pxyl-rnpA* cells grown in the presence or absence of inducer, probed with oligonucleotides targeting different ORFs of the operon (indicated below panel). Colored dots correspond to the colors of the transcripts shown in panel A.

### A determinant for down-regulation of the *rimM* operon is located within the *ylqD* ORF

To further narrow down which ORF was responsible for down-regulation of *rimM* operon expression, we sub-cloned the *ylqD-rimM* or *rimM*-only parts of the operon under control of a *Pspac* promoter, rendered constitutive by deleting the *lac* operator (Pspac(con); Table S3), with an artificial terminator hairpin to provide a defined 3’ end. The constructs were integrated into the chromosome at the *amyE* locus and levels of the ectopic transcript were analyzed by Northern blot in RNase P-depleted cells using a probe specific for *rimM*. The steady state levels of the synthetic *ylqD-rimM* transcript were down-regulated in response to RNase P depletion, albeit not as dramatically as the native operon (1.3-vs 2.3-fold), suggesting that a determinant involved in down-regulation is still included in this shorter construct (Figure 5A and B). Two degradation intermediates (~0.5 and ~0.4 kb in size) of the *ylqD-rimM* transcript also accumulated, suggesting that this transcript is cleaved twice under conditions of RnpB-depletion. It is possible that the weaker effect of RnpB-depletion the full-length transcript and the accumulation of visible degradation intermediates is explained by the presence of a stabilizing terminator hairpin at the 3’ end of each of these species that is not present in the native mRNA. Intriguingly, the construct containing only the *rimM* ORF (with the same 3’ terminator) was up-regulated in response to RNase P depletion (Figure 5C and D). In combination, these results suggest that the region responsible for post-transcriptional down-regulation of *rimM*-containing transcripts upon depletion of RNase P is primarily located within the *ylqD* ORF.

**Figure 5.**
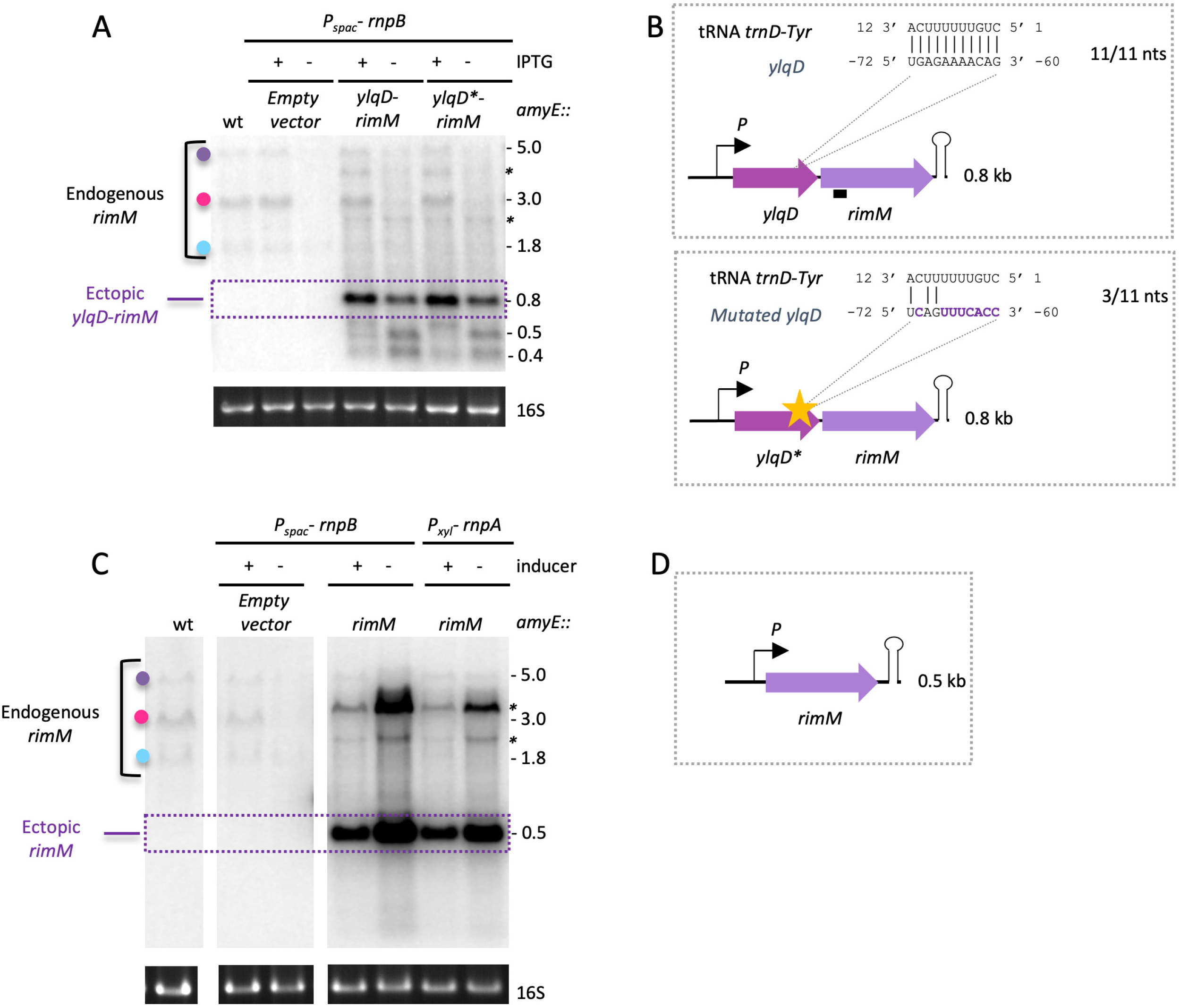
A determinant for down regulation of the *rimM* operon in response to tRNA maturase depletion is located within the *ylqD* ORF. (A) Northern blot of total RNA from *Pspac-rnpB* cells isolated in the presence or absence of IPTG showing the effect of RnpB-depletion on expression of an ectopic *ylqD-rimM* short operon containing a wt or mutated (*) potential target sequence for *trnD-Tyr* pre-tRNA within the *ylqD* ORF. Colored dots identifying endogenous *rimM* transcripts follow the same code as in Figure 4. (B) Schematic of ectopic *ylqD-rimM* constructs placed under control of the constitutive promoter (P) used in panel A. The zoom in shows the complementarity to the *trnD-Tyr* pre-tRNA and its disruption in the *ylqD*-rimM* mutant construct. Coordinates are relative to the start codon of *rimM*. (C) Northern blot showing the effect of RNase P depletion (*rnpA* or *rnpB*) on expression of an ectopic *rimM*-only construct. (D) Schematic of ectopic *rimM* construct placed under control of the constitutive promoter (P) used in panel C. Slower migrating bands (marked with an asterisk) are likely due to read-through of the terminator in the ectopic construct. 16S rRNA levels (ethidium bromide stained) are shown as a loading control.

Since unprocessed tRNAs accumulate in RNase P and RNase Z depleted cells, we wondered whether they could act as potential post-transcriptional regulators of target mRNAs by base pairing to their targets *via* their single stranded 5’ and 3’ extensions. Using TargetRNA2 (12), a prediction program used for identifying targets of small RNAs (sRNAs) in bacteria, we identified an 11-nt region within the *ylqD* ORF that could potentially base-pair with the 5’ immature extension of unprocessed *trnD-Tyr* tRNA (Figure 5B). To test whether this sequence was involved in down-regulation of the *ylqD-rimM* construct in cells depleted for RNase P, we weakened the putative base pairing interaction by introducing mutations in the *ylqD* mRNA sequence (while maintaining the YlqD amino acid sequence as much as possible) (Figure 5B). The mutant *ylqD*-rimM* construct was down-regulated and processed similarly to the wt under conditions of RNase P depletion, suggesting that 5’ extended *trnD-Tyr* does not act as a post-transcriptional regulator of this operon. For the moment, the sequence element(s) within *ylqD* responsible for down-regulation of the *rimM* operon under conditions of tRNA maturase depletion remain(s) unknown.

### Down-regulation of *rimM* expression under physiological conditions resulting in reduced *rnpA* expression is independent of immature tRNA accumulation

We next asked whether the down-regulation of the *rimM operon* we observed during RNase P depletion, would also occur in physiological conditions where RNase P expression is reduced. The level of expression of the *rnpB* RNA is relatively constant in tiling array experiments in over a hundred conditions tested, whereas *rnpA* mRNA levels decrease upon ethanol addition and during stationary phase in both complex and minimal media (Figure 6A) (10). We confirmed that *rnpA* RNA expression was reduced to levels below detection in these three conditions in comparison with exponential growth in the respective medium, by Northern blot (Figure 6B). Ethanol treatment did not affect *rnpB* RNA levels; however, they were reduced during stationary phase in both minimal and complex medium, in contrast to the tiling array data. The expression of *rimM* varied similarly to *rnpA* in the conditions tested (Figure 6B).

**Figure 6.**
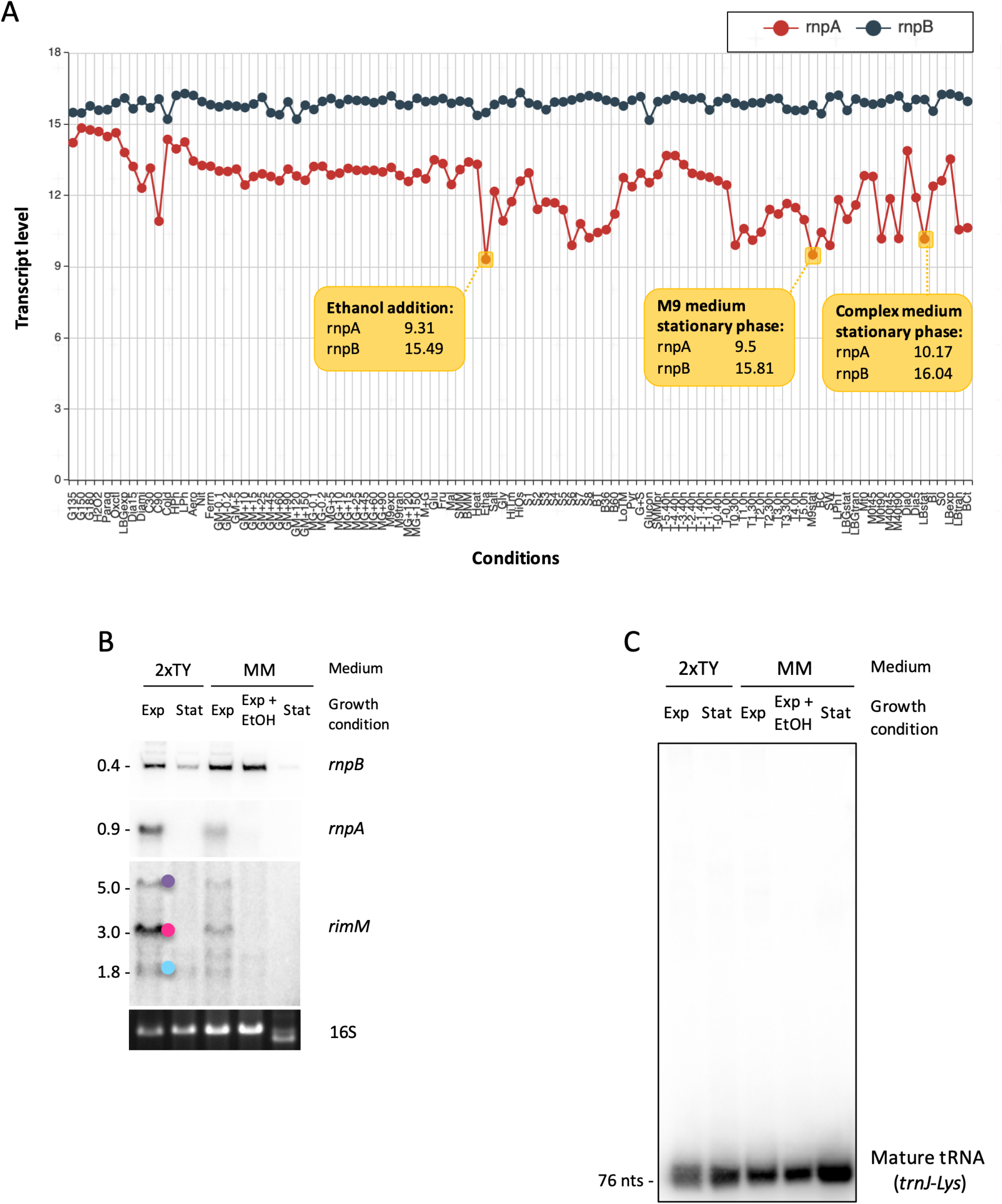
Ethanol stress and stationary phase affect *rnpA* and *rnpB* expression without causing tRNA processing defects. (A) *rnpB* (black) and *rnpA* (red) transcript levels over 100 different growth conditions (from ref. (10)). The three conditions indicated in yellow (ethanol stress and stationary phase in complex and minimal medium) result in reduced *rnpA* RNA levels. For each condition, *rnpA* and *rnpB* RNA levels (log2) are indicated in the yellow box. (B) Northern blot comparing *rnpB* (first panel, acrylamide gel because of small size, 401 nts), *rnpA* and *rimM* (second and third panel, agarose gel) RNA levels, after ethanol addition (EtOH) or during exponential (Exp) or stationary (Stat) phase in minimum (MM) or complex (2xTY) medium. Colored dots identifying *rimM* transcripts follow the same code as in Figure 4. Note that the 16S rRNA (loading control) is beginning to be degraded in MM stationary phase. (C) Northern blot probed for *trnJ-Lys* (acrylamide gel) showing no pre-tRNA accumulation in the different conditions tested. See Figure S1 of ref. (8) for accumulation of *trnJ-Lys* precursors under conditions of RnpA and RnpB depletion.

We next asked whether tRNA maturation was affected in stationary phase or upon addition of ethanol using a probe for *trnJ-Lys* tRNA. Surprisingly, despite the decreased levels of the *rnpA* mRNA in all three conditions, and *rnpB* in stationary phase, we did not observe an accumulation of pre-tRNAs (Figure 6C). It is possible that very few new tRNA molecules are synthesized under these conditions and/or that the remaining cellular RNase P activity provided by the stable RnpA protein and RnpB RNA is sufficient to ensure the processing of any that are transcribed. In either case, these experiments suggest that the down-regulation of *rimM* expression that accompanies the decrease in *rnpA* and *rnpB* expression in stationary phase or ethanol stress is more related to growth arrest than an accumulation of immature tRNAs.

### Down-regulation of *rimM* in RNase P depletion strains depends partially on (p)ppGpp production

In bacteria, both stationary phase and ethanol stress are associated with increased production of (p)ppGpp, hyperphosphorylated guanosine derivatives that are known to globally reprogram transcription (13, 14). Considering that tRNA maturase-depleted cells also trigger a RelA-dependent production of (p)ppGpp (8), we asked whether *rimM* down-regulation in these cells was dependent on (p)ppGpp production, by measuring *rimM* expression in (p)ppGpp^0^ strains depleted for RnpA or RnpB. The (p)ppGpp^0^ strain lacks the three genes encoding (p)ppGpp synthesizing enzymes in *B. subtilis* (*yjbM, ywaC* and *relA*) (15). If (p)ppGpp were the key mediator, we would expect that the effect of RNase P depletion on *rimM* expression to be reduced or abolished in the (p)ppGpp^0^ background. Rather than simply abolishing the effect, *rimM* transcripts actually showed higher levels in the tRNA maturase-depleted (p)ppGpp^0^ strains compared to the RnpA or RnpB-depleted strains capable of making (p)ppGpp (Figure 7A), suggesting that (p)ppGpp has an independent repressive effect on *rimM* mRNA levels.

**Figure 7.**
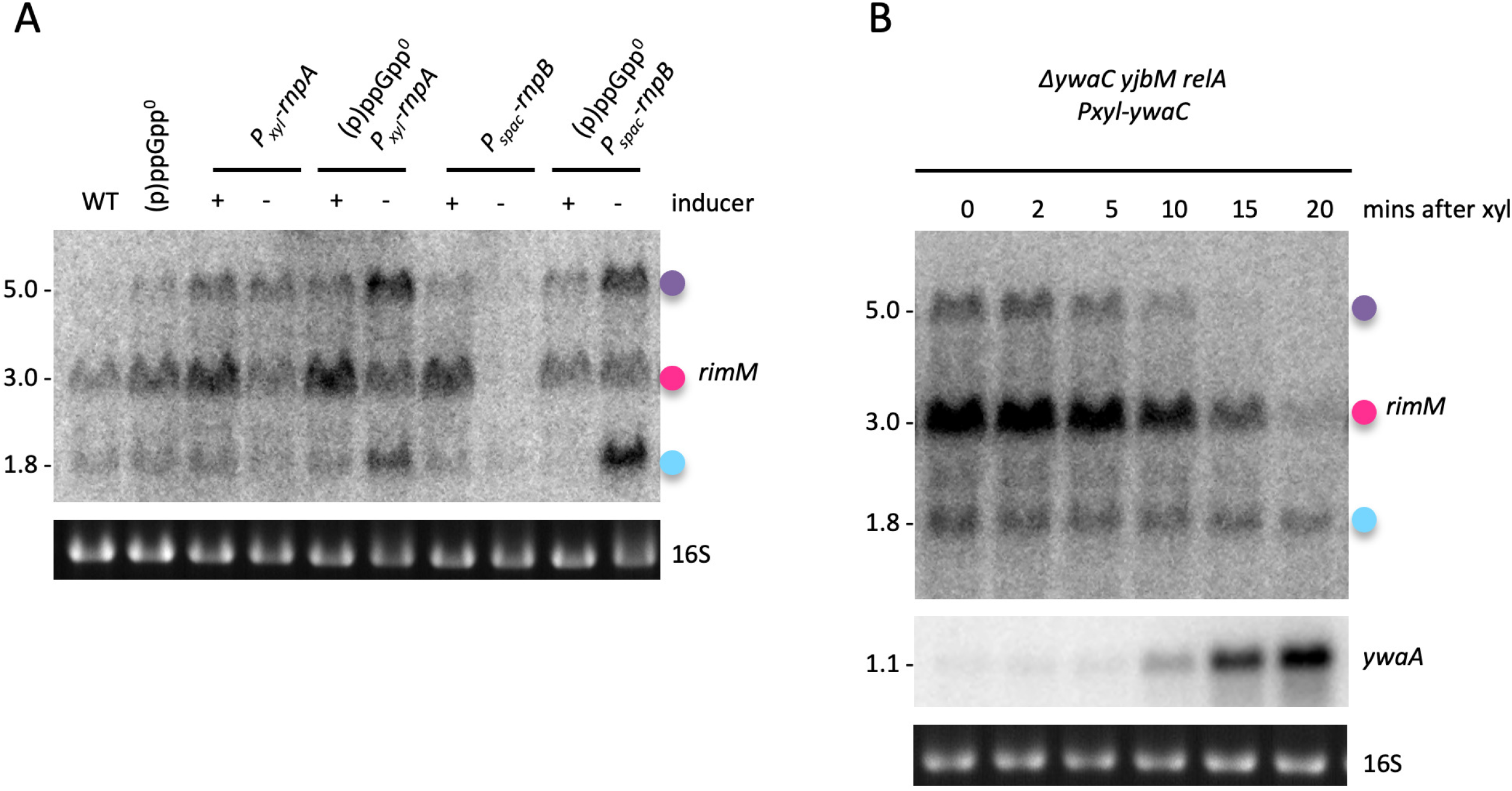
Influence of (p)ppGpp on *rimM* expression. (A) Northern blot comparing the effect of RNase P depletion (RnpB or RnpA) on *rimM* expression in a wt and (p)ppGpp^0^ background. (B) Northern blot of *rimM* expression after induction of (p)ppGpp production using a xylose inducible *ywaC* in a *ΔyjbM ywaC relA* ((p)ppGpp^0^) background. Colored dots identifying *rimM* transcripts follow the same code as in Figure 4. 16S rRNA levels (ethidium bromide stained) are shown as a loading control. Series of blots where a single loading control is shown, were stripped and reprobed. Note that this Northern blot was generated by reprobing a membrane previously used in ref. (8), with permission to re-use the *ywaA* control panel granted by the publisher.

We thus assessed whether the alarmone (p)ppGpp could down-regulate *rimM* expression in the absence of a tRNA processing defect using an engineered strain that allows us to produce (p)ppGpp in the absence of immature tRNA accumulation or nutrient starvation (8). This (p)ppGpp^+^ strain consists of an ectopic copy of the *ywaC* gene placed under the control of a *Pxyl* promoter in the (p)ppGpp^0^ strain background. We used derepression of the CodY-regulated *ywaA* mRNA as a proxy to follow the increase in (p)ppGpp levels in this strain *in vivo* (Figure 7B) (8). We observed that (p)ppGpp induction alone had no effect on the small (Y/P_2_-T_2_; cyan dot) *rimM* transcript, whereas the two larger species (Y/P_2_-T_3_ and P_1_-T_3_, pink and purple dots, respectively) were down-regulated as the expression of the (p)ppGpp reporter *ywaA* increased (Figure 7B). Although (p)ppGpp production alone recapitulated what was seen in RnpB-depleted cells, the fact that only the two larger transcripts behaved as expected from the results obtained in the RNase P-depleted ppGpp^0^ strain (Figure 7A), suggests that regulation of the smallest transcript (Y/P_2_-T_2_) is more complex than simple transcriptional repression by (p)ppGpp.

The fact that the Y/P_2_-T_2_ *rimM* transcript could be down-regulated independently of (p)ppGpp production led us to investigate the possibility of a further layer of regulation where the growth slow-down in tRNA maturase-depleted cells would also affect *rimM* expression by a mechanism independent of alarmone levels. To test this, we sought to reproduce the growth rate defect by depleting for an unrelated essential enzyme. We therefore performed Northern blot analysis on total RNA extracted from both RNase III (*rnc*) depletion and deletion strains. The double-strand specific endoribonuclease RNase III is essential in *B. subtilis* because it is required to silence expression of foreign toxin genes of two prophages (Skin and SPβ) (16). Whereas depletion of RNase III in a WT background leads to growth arrest, the *rnc* gene can be deleted in a strain lacking the two prophages without a marked effect on growth rate. The RNase III depleted strain showed only a very limited derepression of the CodY regulon in comparison with tRNA maturase-depleted strains (Figure 8A) and did not accumulate visible amounts of (p)ppGpp on thin-layer chromatography (TLC) (Figure 8B). This validates the use of RNase III depletion strains to examine the effect of growth rate on *rimM* expression and to distinguish this from the effect of accumulating high levels of (p)ppGpp. While RNase III deletion had no effect on *rimM* expression, all three *rimM-*containing transcripts were strongly down-regulated during RNase III depletion (Figure 8C), confirming that growth rate also plays a major role in the regulation of *rimM* expression, independently of (p)ppGpp and tRNA maturase depletion.

**Figure 8.**
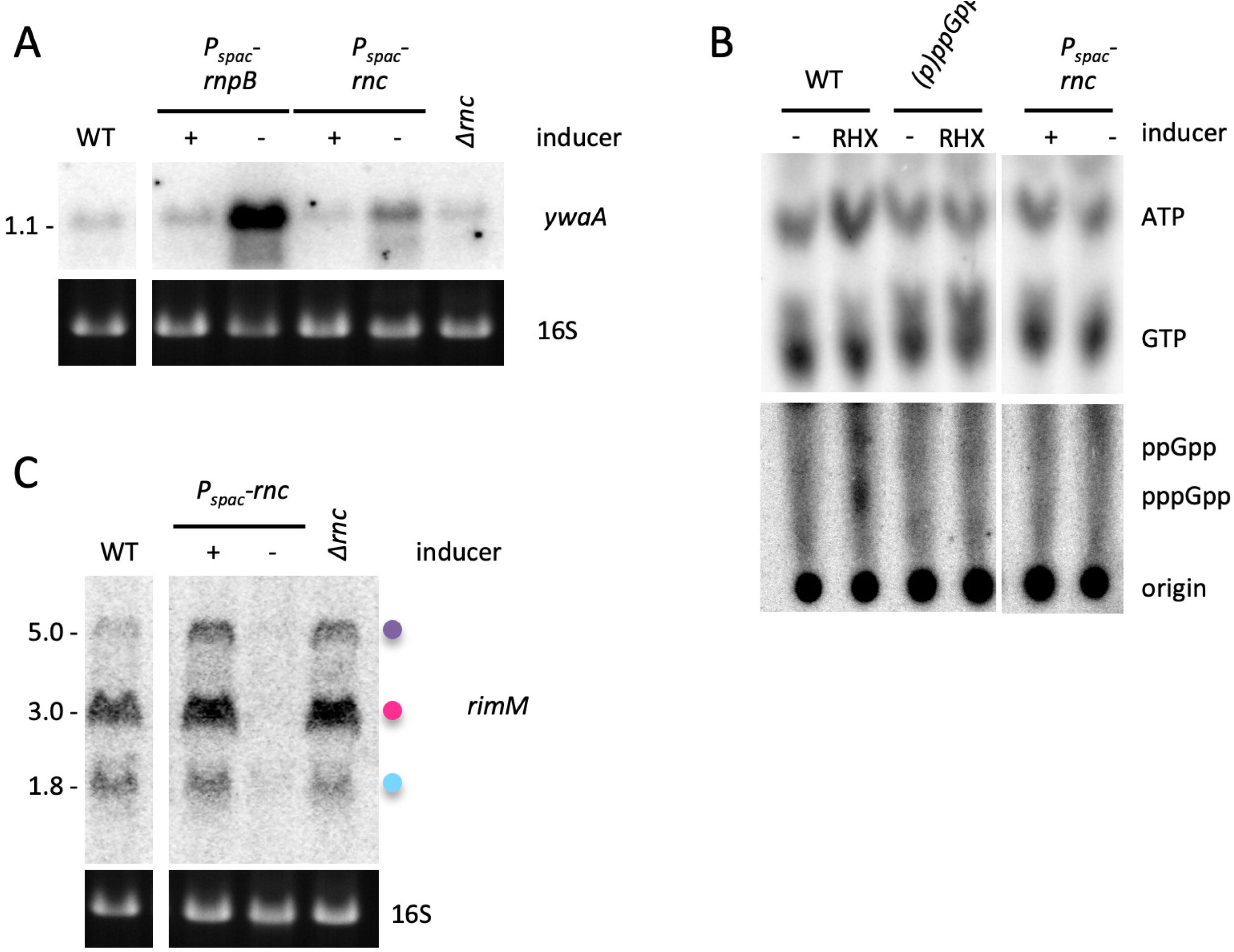
Expression of the *rimM* operon is regulated by growth rate independent of (p)ppGpp. (A) Northern blot comparing the effect of RNase III deletion, and RNase P or RNase III depletion on derepression of the CodY-regulated *ywaA mRNA*. (B) RNase III-depleted cells do not accumulate large amounts of (p)ppGpp compared to tRNA maturase depleted ones. Thin-layer chromatography (TLC) analysis of ^32^P-labeled nucleotides extracted from RNase III-depleted cells (*rnc*). Arginine hydroxamate (RHX; 250mg/mL) was added to wt and (p)ppGpp^0^ strains as positive and negative controls. The top and bottom halves are exposed for different times. Note that this is a re-crop of an image previously published in ref. (8); the first four (control) lanes are reshown to show the migration position of (p)ppGpp, with permission from the publisher. (C) Northern blot comparing the effect of RNase III depletion or deletion on *rimM* expression. Colored dots identifying *rimM* transcripts follow the same code as in Figure 4. 16S rRNA levels (ethidium bromide stained) are shown as a loading control.

## Discussion

This study began with the observation that depletion of tRNA maturase enzymes in *B. subtilis* led to a defect in 30S ribosome subunit assembly, which we showed was in part due to an accumulation of (p)ppGpp and an inhibition of the activity of 30S assembly GTPases (8). In our early attempts to understand the mechanism underlying this phenomenon, we studied the expression of several 30S assembly factors and discovered that most were either up or down-regulated at the mRNA level upon depletion of either RNase P or RNase Z.

We initially focused on the *rimM* mRNA because the 30S ribosome assembly defect observed in tRNA maturase depletion mutants was very similar to that seen in a *△rimM* strain. Although we later showed that ectopic *rimM* expression could not correct the assembly defect in the *rnpB*-depleted strain (8), we were nonetheless curious about how *rimM* expression was affected by the decrease in the levels of mature tRNAs. Together, our data (summarized in Figure S4) indicate that the down-regulation of *rimM* transcript levels in tRNA maturase-depleted cells is the result of a complex mixture of transcriptional and post-transcriptional mechanisms, caused by a combination of effects mediated by a reduction in growth rate, (p)ppGpp production and a translational defect due to lack of functional tRNAs, with each layer of regulation capable of functioning independently of the others and affecting the three *rimM* transcripts distinctly at different levels of severity.

We hypothesized that the accumulation of immature tRNAs during tRNA maturase depletion increases ribosome stalling. Stalled ribosomes are known to affect mRNA decay in bacteria (17) and a tRNA loss of function mutation leading to pre-tRNA processing defects was reported to induce ribosome stalling in mice (9). In agreement with our hypothesis, we observed that treatment with the translation elongation inhibitor chloramphenicol at MIC concentrations recapitulates the effects of tRNA maturase depletion on the mRNA levels of several different assembly factor mRNAs tested, four of which were up-regulated (*era, yqeH, ydaF and yjcK*) and two down-regulated (*rimM* and *cpgA*). Interestingly, Cm treatment at sub-inhibitory concentrations did not impact cofactor mRNA levels in the same way, with low Cm concentrations initially having transitory up-effects that were then reversed at longer incubation times. One possibility is that short ribosome stalls transiently block access to cleavage sites by housekeeping RNases such as RNase Y, resulting in mRNA stabilization, while prolonged stalling could lead to mRNA destabilisation by an enzyme such as Rae1, proposed to enter the A-site of stalled ribosomes (2), or by leaving large stretches of mRNA unoccupied by ribosomes and vulnerable to cleavage by canonical degradation pathways.

Another notable difference between the two conditions is that the stringent response is induced by Cm at MIC, as evidenced by the increase in expression the *ilvA* mRNA from the CodY regulon (Figure S5), which could also contribute the increased severity of the response to higher Cm concentrations. Activation of the stringent response in Cm-treated *B. subtilis* was also previously observed by (18), consistent with our results. This is a marked difference from *E. coli*, where Cm is a known inhibitor of stringent response induction (19, 20). The mechanism still remains elusive in both cases.

Beyond their canonical role in protein synthesis, tRNAs have been implicated in the regulation of several biological processes (for review, see (21, 22)). A new class of small non-coding RNAs has emerged recently called tRNA-derived fragments (tRFs) or tRNA-derived small RNAs, whose biological roles are not yet well understood (23). Different types of tRFs differ in the cleavage position of the mature or precursor tRNA transcript. They have been particularly studied in humans, where they have been shown to be involved in regulation of a variety of cellular processes, including global translation, cellular proliferation, apoptosis and epigenetic inheritance (24). Interestingly, a 3’-tRF in human cells plays an essential role in fine-tuning ribosome biogenesis under normal physiological conditions by post-transcriptionally regulating translation of at least two r-protein mRNAs (25). Although tRFs have not yet been identified in *B. subtilis*, we asked whether pre-tRNAs could bind certain assembly factor mRNAs *via* their 5’ or 3’ extensions and cause some of the post-transcriptional effects observed in the tRNA maturase depletion strains. tRFs with 5’ or 3’ extensions (pre-tRFs) could similarly behave as a new pool of potential regulatory sRNAs. Although the potential base-pairing we identified between the 5’ extension of trnD-Tyr and the *rimM* transcripts does not seem to play a role in the down-regulation of *rimM* expression, this doesn’t preclude the possibility that other pre-tRNAs or pre-tRFs could be involved in post-transcriptional regulatory events in *B. subtilis*.

A recent study in *E. coli* showed that the abundance of 46% of transcripts were affected in a strain where the protein moiety of RNase P was heat denatured (26). The observation that the addition of chloramphenicol mimicked the effect of tRNA maturase depletion in *B. subtilis* for the upregulated mRNAs, and that down-regulation was the net result of a mixture of translational and transcriptional effects, suggests that the effects seen on the *E. coli* transcriptome may substantially be the result of ribosome stalling on mRNAs due to lack of functional tRNAs, with differential impacts (up, down or neutral) on individual mRNA stabilities or transcription levels. More detailed studies are required to untangle these effects on a global level in both organisms.

## Acknowledgements

This work was supported by funds from the CNRS (UMR 8261), Université Paris Cité, the Agence Nationale de la Recherche (ARNr-QC). This work has been published under the framework of Equipex (Cacsice) and LABEX (Dynamo) programs that benefit from a state funding managed by the French National Research Agency as part of the Investments for the Future program. We thank lab members for helpful discussion.

## Materials and Methods

### Strains and culture conditions

All *B. subtilis* strains used were derived from our laboratory strain SSB1002, a W168 *trp^+^* prototrophic strain. Strains are listed in Table S1 and details of strains and plasmid constructs are provided in Table S2 and S3, respectively. Oligonucleotides used are listed in Table S4.

Unless stated otherwise, *B. subtilis* strains were grown in 2xYT liquid medium (1.6% peptone, 1% yeast extract, 1% NaCl) at 200 rpm at 37°C in ≤ 1/10 volume of the flask to ensure proper aeration. Overnight precultures were grown in presence of appropriate antibiotics and inducer (1mM IPTG or 2% xylose), in the case of depletion strains. Experimental cultures were grown in the absence of antibiotics, except where stated. For depletion strains, overnight induced cultures were washed three times with pre-warmed 2xYT medium and inoculated at OD_600_ between 0.02 and 0.2, depending on the strain, in fresh medium with or without inducer. Generally, induced cells were harvested for RNA preparation around OD_600_ = 0.6 and cells grown in the absence of the inducer were followed until they reach a plateau before being harvested. Inoculation and depletion conditions were determined empirically for each strain such that the depleted cells were harvested between OD_600_ = 0.3 and 0.7. For RnpA depletion, cultures were inoculated at OD_600_ = 0.05 in presence of 2% xylose (inducer) or 2% glucose to tighten repression of the *Pxyl* promoter, which typically led to a growth arrest (plateau) around OD_600_ = 0.6. For *rnz* and *rnpB* depletion strains, cultures were inoculated in presence or in absence of 1mM IPTG at OD_600_ = 0.05 and OD_600_ = 0.2, respectively. RNase Z and RnpB depleted cells typically plateau around OD_600_ = 0.6 and OD_600_= 0.3, respectively.

For rifampicin experiments, *B. subtilis* strains were grown in 2xTY at 37°C with shaking as described above. At OD_600nm_ = 0.6 (or less for some depletion strains), rifampicin was added to a final concentration of 150 μg/mL in order to block new RNA synthesis. Samples were collected at different time points (e. g. 0, 2, 5, 10, 15 and 20 minutes) by mixing the cells with frozen 10 mM sodium azide (200 μL for 1.3 mL culture). Samples were vortexed until the sodium azide thawed, cells were pelleted by centrifugation at 4°C and the pellet was conserved at −20°C until RNA extraction.

To mimic amino acid starvation, we depleted charged arginine tRNAs by addition of arginine hydroxamate (RHX) at 250 mg/mL in cultures growing in 2xTY at OD_600_ = 0.3.

To study the effect of translation pausing, we added the translation elongation inhibitor chloramphenicol (Cm) at sub-inhibitory (2.5 μg/ml) or minimal inhibitory concentration (5 μg/ml) to cells growing in 2xYT at OD_600_ = 0.6. Cells were harvested just before Cm addition (t0) and 15, 30 and 60 mins after treatment.

To reproduce some growth conditions from the *B. subtilis* tiling array experiment (10) known to lead to a decrease in *rnpA* expression, ethanol was added to cultures growing in minimal medium (M9 with 0.5 % glucose) at 4% (v/v) around OD_600_ = 0.4 and cells were harvested 10 mins after treatment.

### Plasmid constructs

The *rimM* gene was amplified by PCR using oligo pair (CC2034/CC1986) and cloned between the BamHI and XhoI sites of the integrative pHM2-Pspac(con) vector (Table S3). The bicistronic *ylqD-rimM* construct was amplified by PCR using oligo pair (CC1985/CC1986) and cloned between the BamHI and SalI sites of pHM2-Pspac(con). The mutated construct *ylqD*-rimM* was obtained by two-fragment overlapping PCR. The upstream fragment was amplified with the forward primer CC1985 and the reverse primer CC2012 and the downstream fragment with the forward primer CC2011 and the reverse primer CC1986. The overlapping fragments were reamplified using oligo pair CC1985/CC1986 and cloned between the BamHI and SalI sites of pHM2-Pspac(con). The integrative plasmids were linearized with XbaI before transformation, for integration into the *amyE* locus of the *B. subtilis* chromosome.

### RNA extraction and Northern blots

RNA extraction was typically performed using the glass beads/phenol protocol (adapted from (27)) on 1 to 8 mL mid-log phase *B. subtilis* cells growing in 2xYT.

To perform Northern blots, 5 μg total RNA were denatured for 5 mins at 95°C in RNA Gel loading dye (Thermo Scientific) before being separated on 1% agarose gels in 1X TBE (native) or on denaturing 5% acrylamide gels in 1X TBE + 7M urea. RNA was transferred from agarose gels to a hybond-N membrane (GE-Healthcare) by capillary transfer for 4 hours minimum in 1X transfer buffer (5X SSC, 0.01M NaOH). For Northerns of acrylamide gels, RNA was electro-transferred at 4°C in 0.5X TBE for 4 hours at 60V or overnight at 12V. RNA was cross-linked to the membrane by UV cross-linking at 120,000 microjoules/cm^2^ using HL-200 Hybrilinker UV-crosslinker (UVP). Probes for Northern blots were usually 25 to 30-nt DNA oligonucleotides radiolabeled on their 5’ end by polynucleotide kinase. The *cpgA* mRNA was detected using a riboprobe using a PCR fragment amplified using oligos CC2200 and CC2201 as template. Membranes were pre-incubated in Ultra-Hyb (Life Technologies) for agarose blots or Roti-Hybri-Quick (Roth) for acrylamide blots for 1 hour and hybridized with radiolabeled probes for a minimum of 4 hours. Pre-incubation, hybridization and wash steps were performed at 42°C in the case of 5’-labeled oligonucleotides or at 68°C for riboprobes. Membranes were quickly rinsed once at room temperature in 2x SSC 0.1% SDS to remove non-hybridized probe before being washed once for 5 mins in the same buffer and then twice for 5 mins in 0.2x SSC 0.1% SDS. Northerns were exposed to PhosphorImager screens (GE Healthcare) and the signal was obtained by scanning with a Typhoon scanner (GE Healthcare) and analyzed by Fiji (ImageJ) software.

### Thin layer chromotagraphy (TLC)

TLC analysis was used to detect radiolabeled (p)ppGpp as described in (8).

---

**Table S1:**
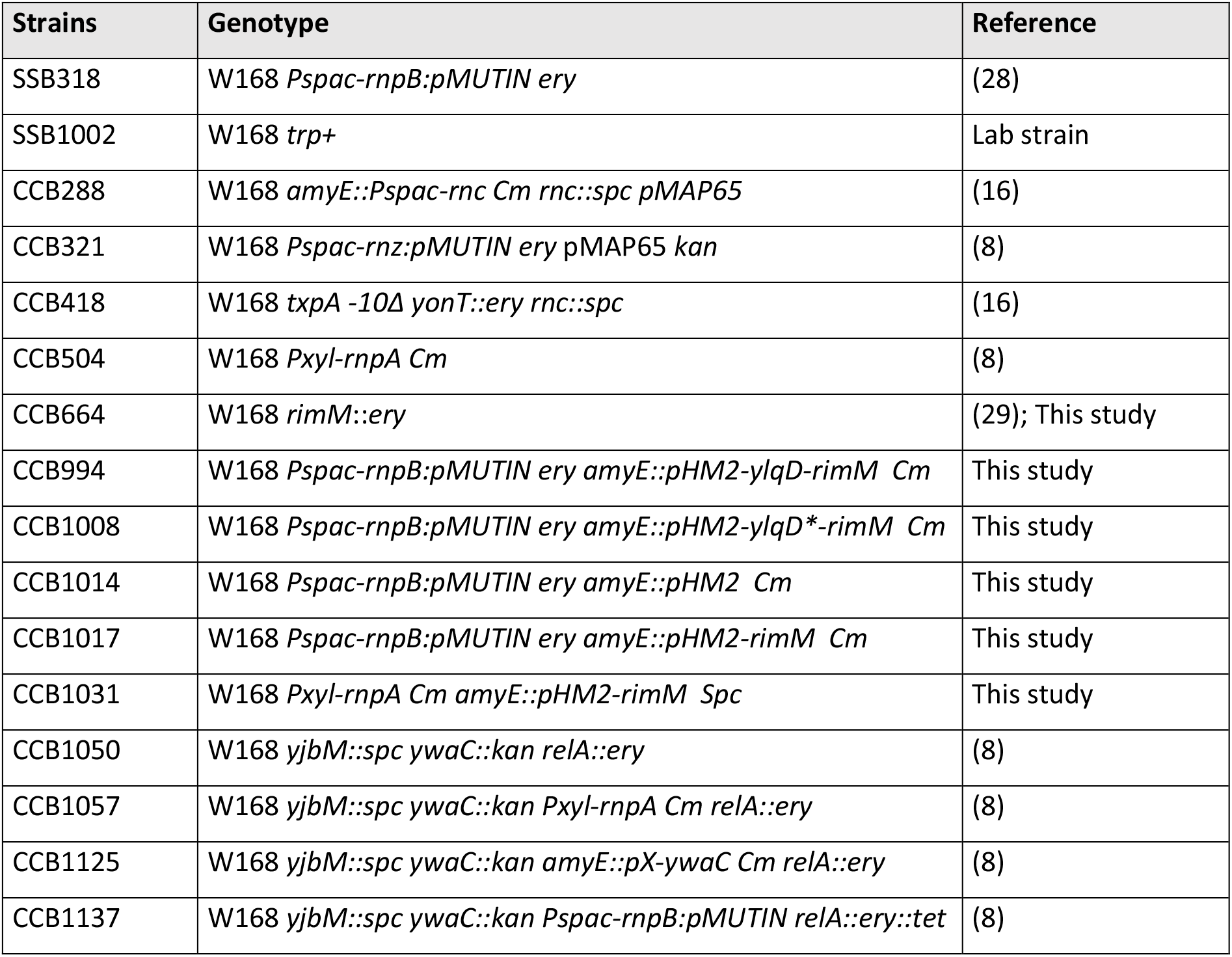
*B. subtilis* strains used in this study.

**Table S2:**
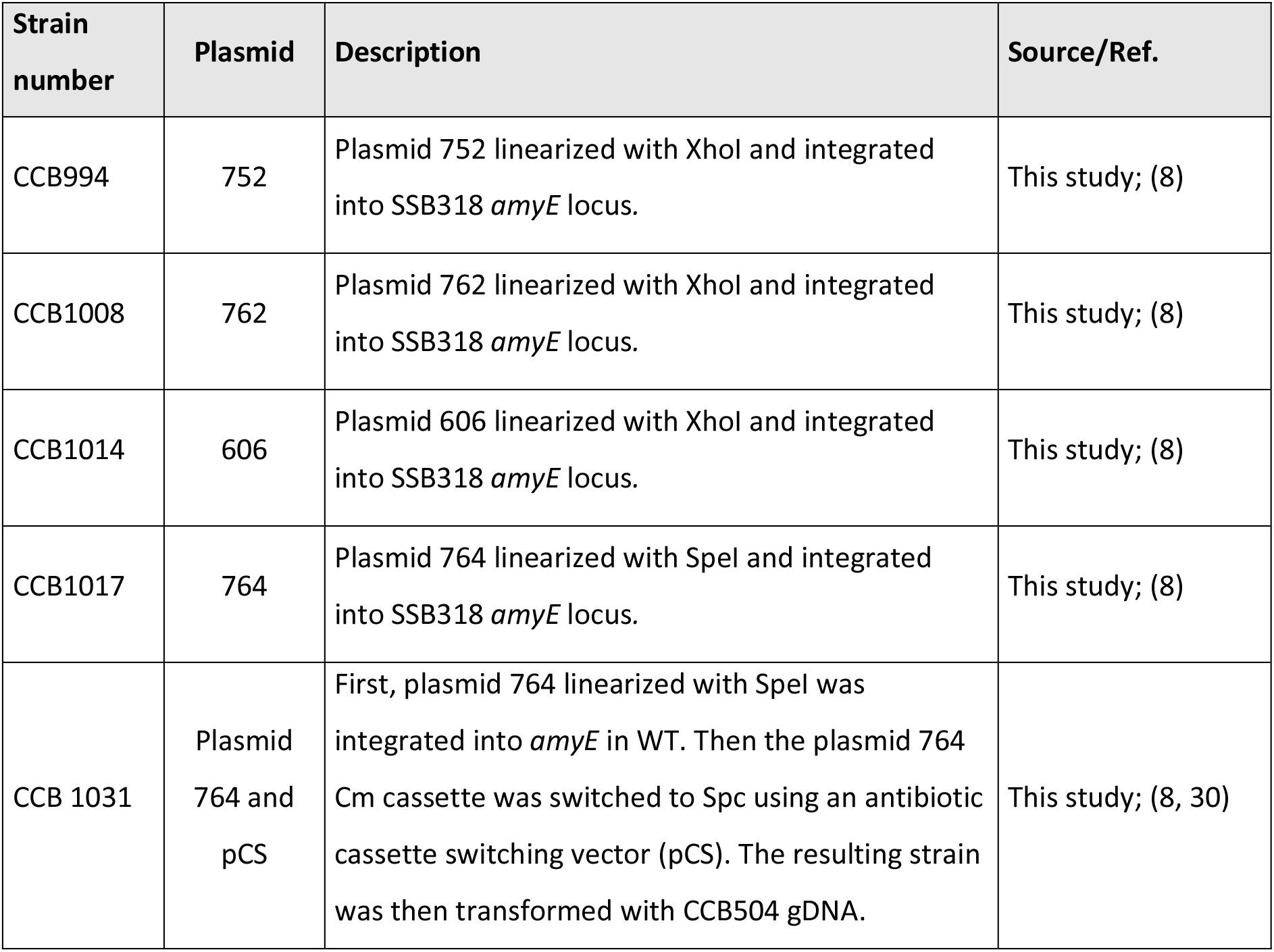
Details of strain construction.

**Table S3:**
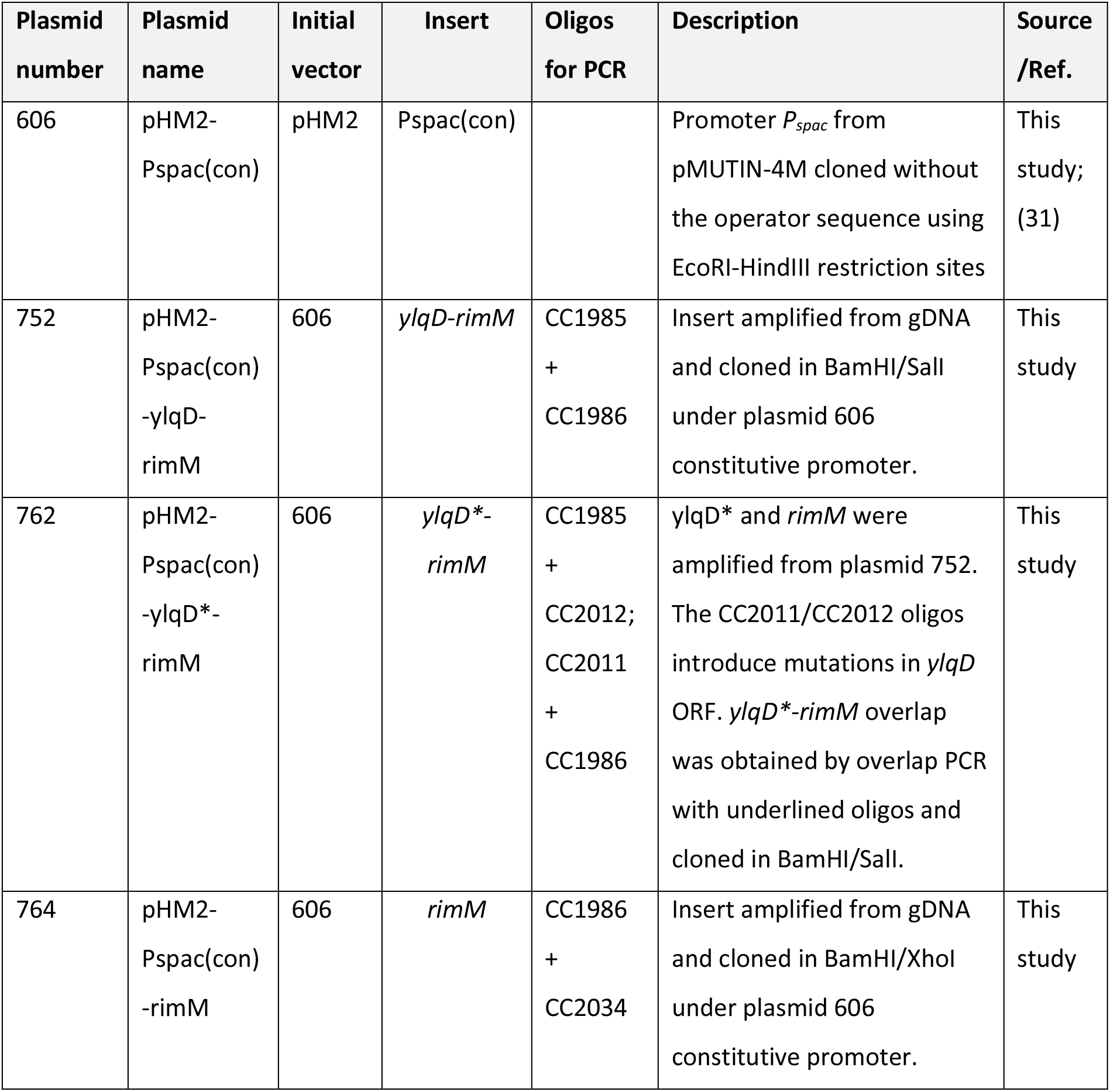
Plasmids constructed for this study.

**Table S4:**
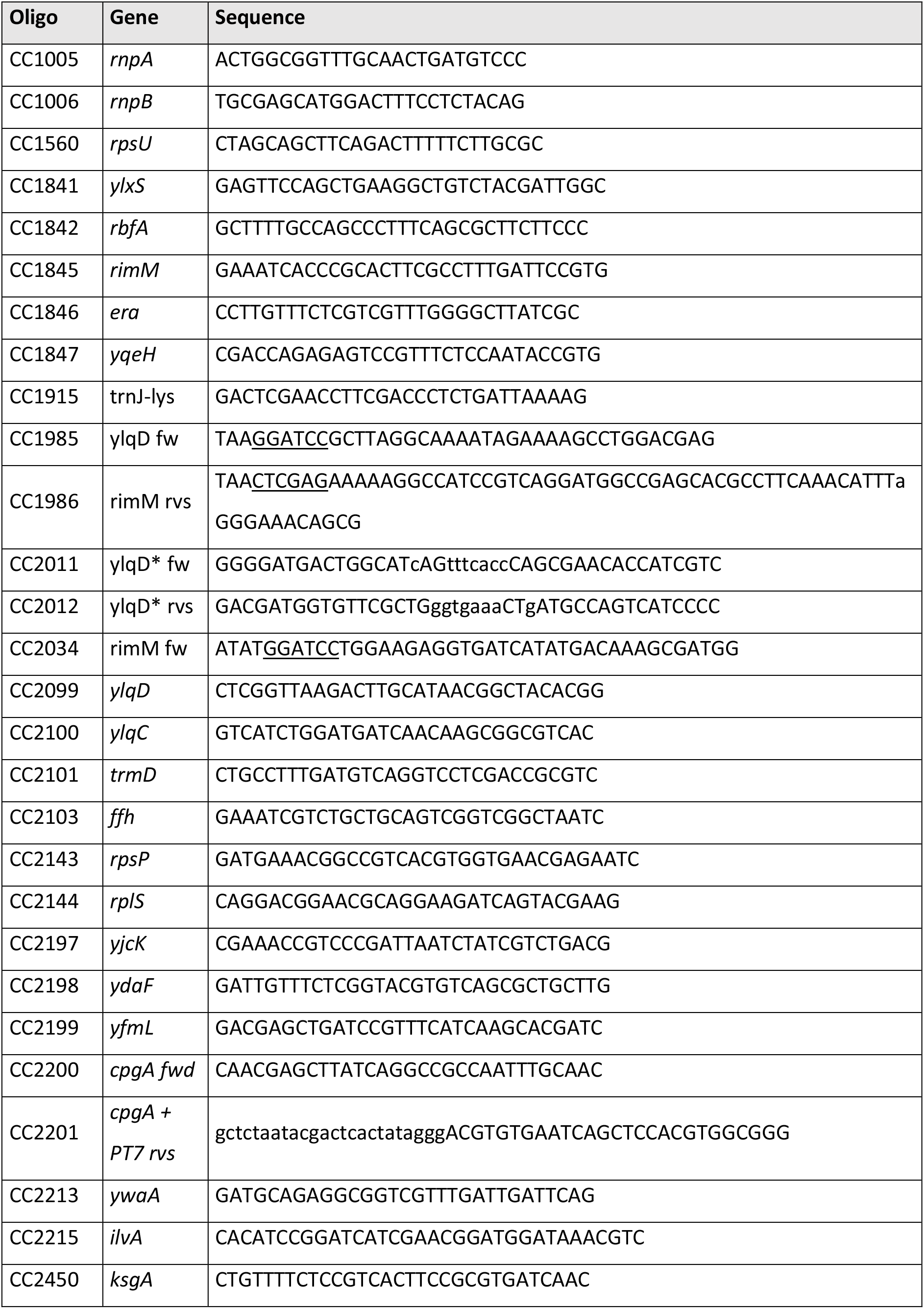
Oligonucleotides used in this study.

**Figure S1:**
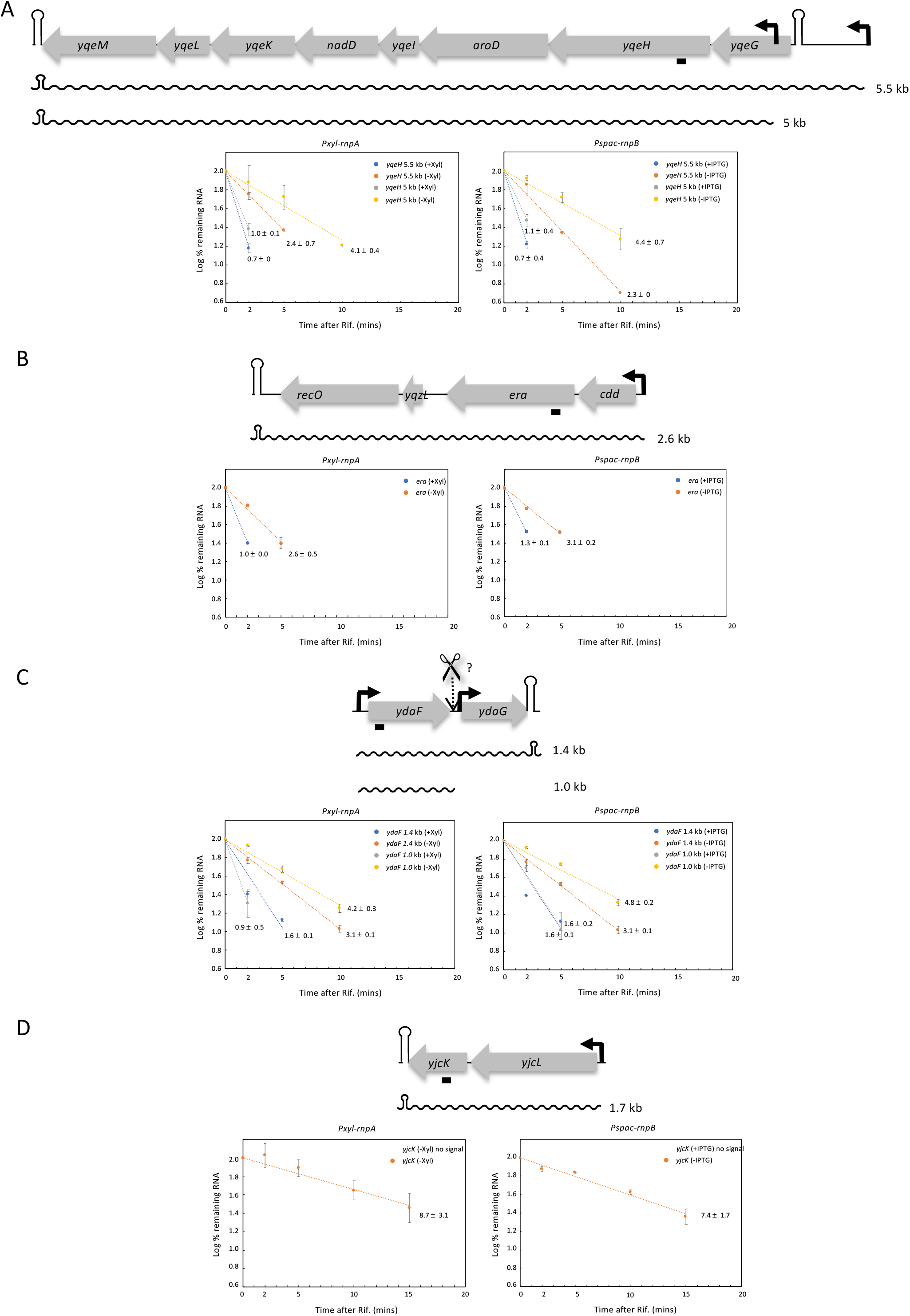
Schematics and RNA decay plots of up-regulated transcripts in RnpA and RnpB-depletion strains. Open reading frames (ORFs; not to scale) are shown as gray arrows and transcripts as wavy lines. Sizes are as indicated. Promoters are represented by black arrows and terminators as hairpins. Suspected endoribonucleolytic cleavage sites are indicated by a scissors symbol with a question mark. Quantifications are from the average of two experiments, with half-lives as indicated.

**Figure S2:**
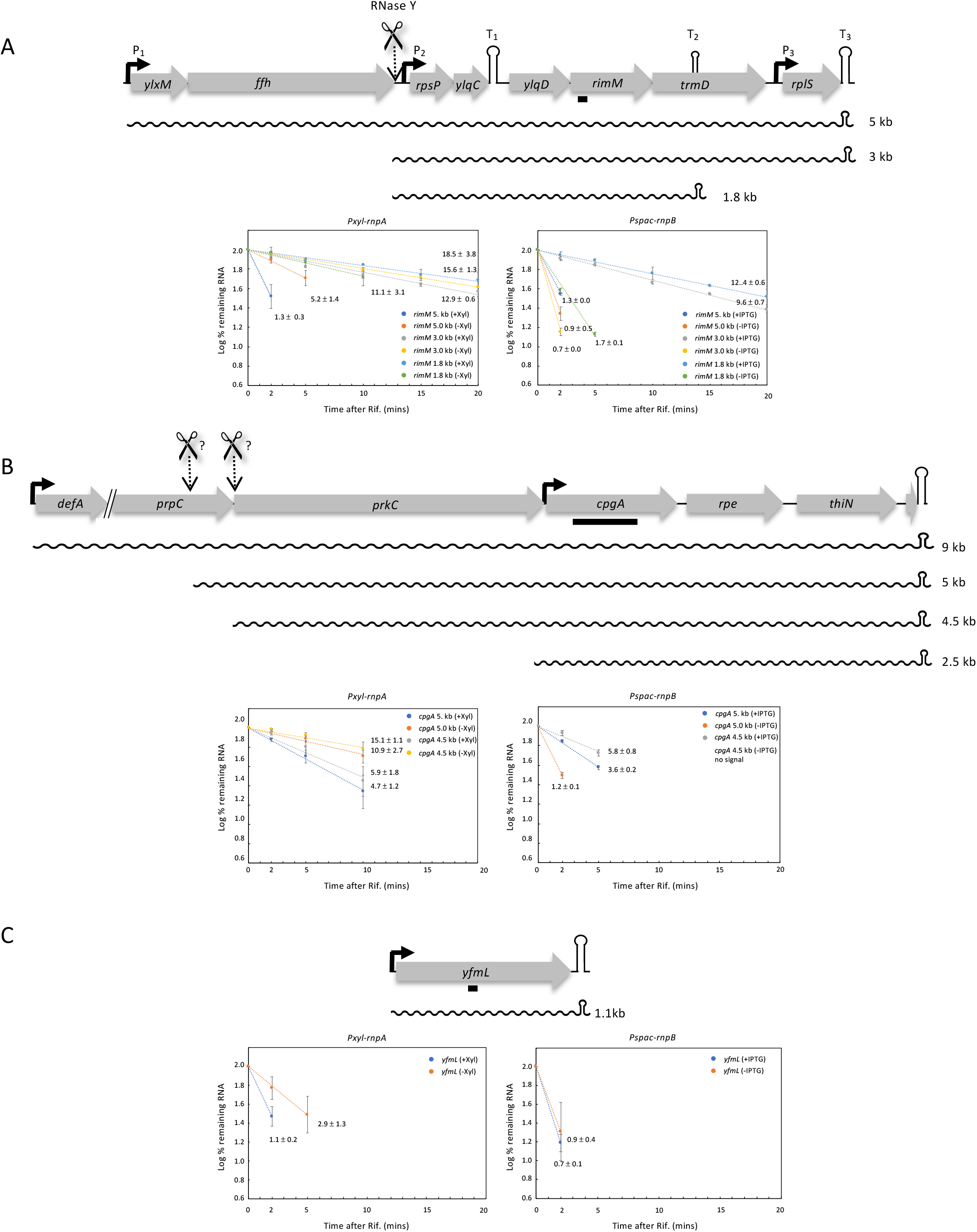
Schematics and RNA decay plots of down-regulated transcripts in RnpA and RnpB-depletion strains. Open reading frames (ORFs; not to scale) are shown as gray arrows and transcripts as wavy lines. Sizes are as indicated. Promoters are represented by black arrows and terminators as hairpins. Suspected endoribonucleolytic cleavage sites are indicated by a scissors symbols with question marks. Quantifications are from the average of two experiments, with half-lives as indicated.

**Figure S3:**
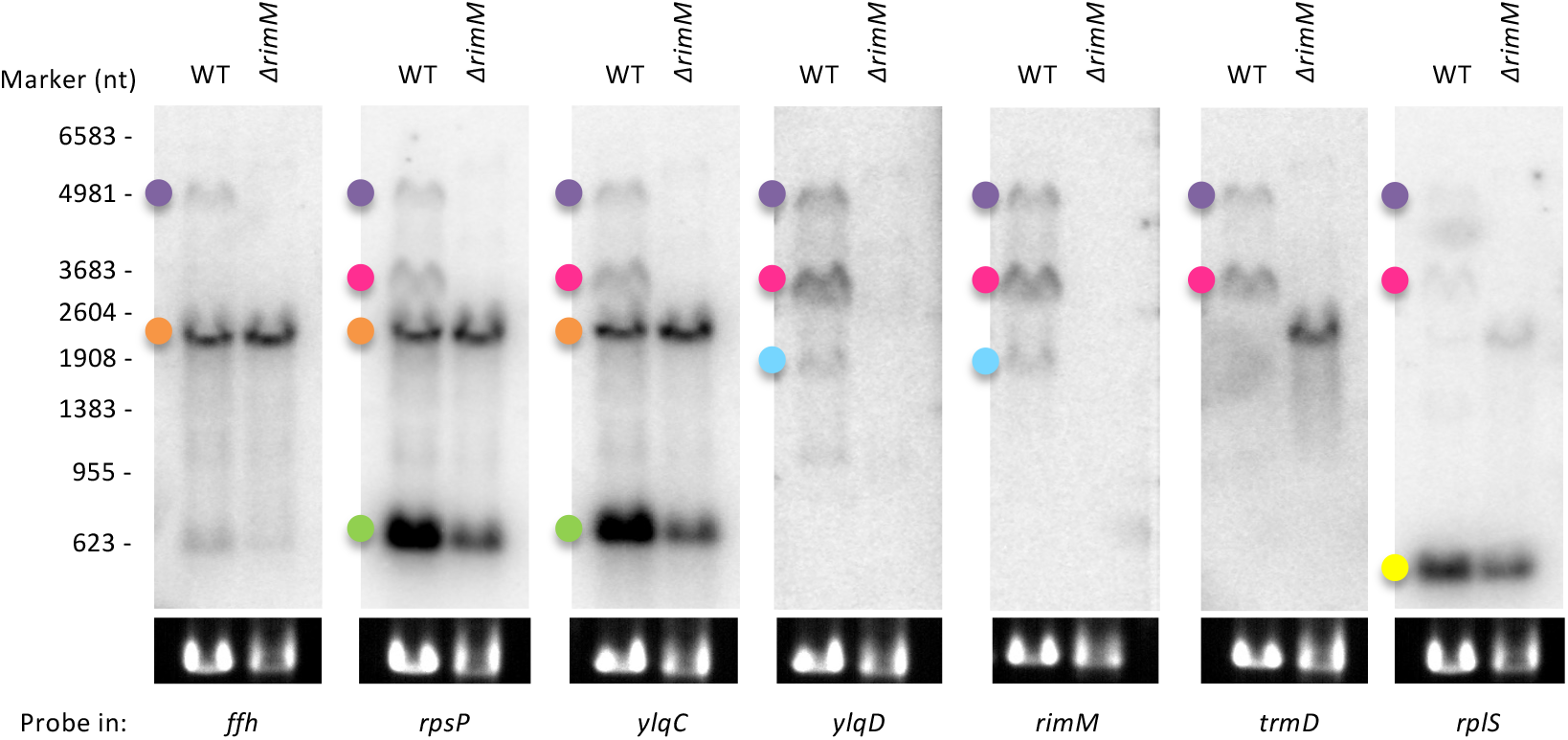
Northern blot analysis to determine the structure of the *rimM* operon. Total RNA from WT or *DrimM* strains was probed with oligonucleotides targeting different ORFs of the *rimM* operon (indicated below each panel). 16S rRNA levels (ethidium bromide stained) are shown as a loading control. The *ffh* and *ylqC* blots were stripped and reprobed for *rplS* and *ylqD*, respectively. Colored dots correspond to the color of transcripts shown in Figure 4A.

**Figure S4.**
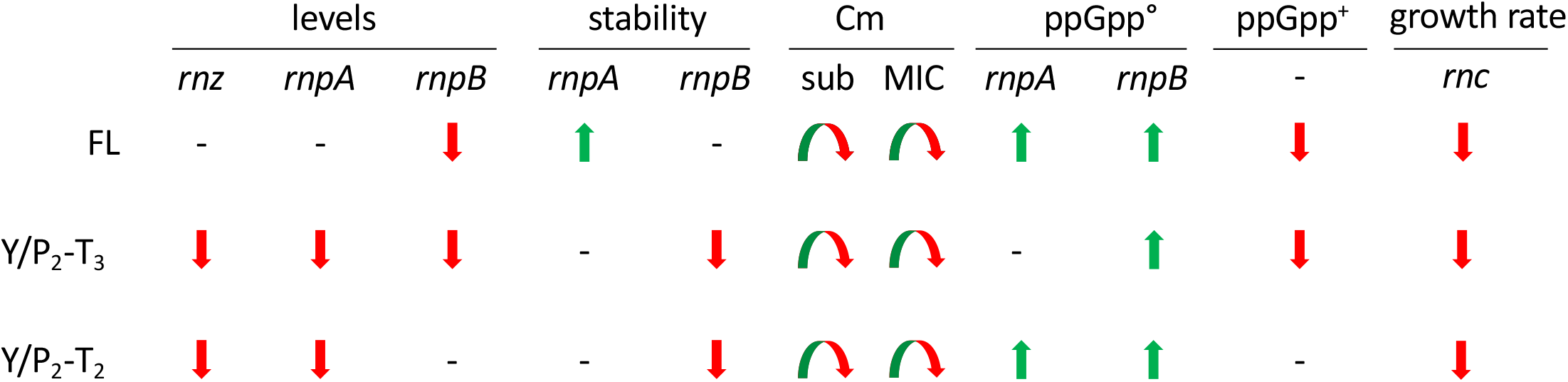
Summary of effects on 3 three *rimM* containing transcripts. Data are taken from Northern blots shown in Figures 1 (levels), 2-3 (stability, Cm), 6A (ppGpp^0^), 6B (ppGpp^+^) and 7 (growth rate). Depleted tRNA maturases are indicated. Transcripts are listed according to size: full-length (FL), RNase Y/P_2_-T_3_ and RNase Y/P_2_-T_2_. Red arrows show down-effects, green arrows show up-effects, (-) no effect. Curved arrow in sub-inhibitory Cm condition (sub) reflects initial increase followed by decrease.

**Figure S5.**
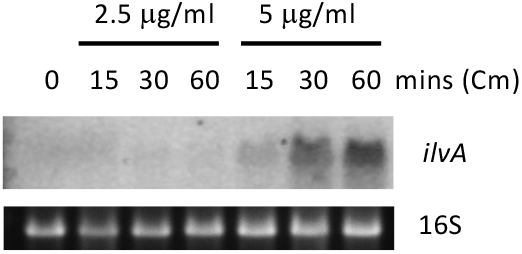
Induction of expression of the CodY regulon member *ilvA* by chloramphenicol. Northern blots of total RNA isolated at different times after addition of 0.5x MIC and MIC of chloramphenicol (Cm). 16S rRNA levels (ethidium bromide stained) are shown as a loading control.

## Notes

### Competing Interest Statement

The authors have declared no competing interest.

### Summary of Updates

Supplementary tables and figures

## References

1. Condon C, Bechhofer DH. 2011. Regulated RNA stability in the Gram positives. Curr Opin Microbiol 14:148–54.

2. Leroy M, Piton J, Gilet L, Pellegrini O, Proux C, Coppee JY, Figaro S, Condon C. 2017. Rae1/YacP, a new endoribonuclease involved in ribosome-dependent mRNA decay in Bacillus subtilis. EMBO J 36:1167–1181.

3. Condon C, Piton J, Braun F. 2018. Distribution of the ribosome associated endonuclease Rae1 and the potential role of conserved amino acids in codon recognition. RNA Biol doi:10.1080/15476286.2018.1454250:1-6.

4. Braun F, Condon C. 2019. RNA processing, p 164–177 In Schmidt TM (ed), Encyclopedia of Microbiology, 4th edition doi:10.1016/B978-0-12-801238-3.02455-7. Elsevier.

5. Wen T, Oussenko IA, Pellegrini O, Bechhofer DH, Condon C. 2005. Ribonuclease PH plays a major role in the exonucleolytic maturation of CCA-containing tRNA precursors in Bacillus subtilis. Nucleic Acids Res 33:3636–43.

6. Pellegrini O, Li de la Sierra-Gallay I, Piton J, Gilet L, Condon C. 2012. Activation of tRNA maturation by downstream uracil residues in B. subtilis. Structure 20:1769–77.

7. Marks J, Kannan K, Roncase EJ, Klepacki D, Kefi A, Orelle C, Vazquez-Laslop N, Mankin AS. 2016. Context-specific inhibition of translation by ribosomal antibiotics targeting the peptidyl transferase center. Proc Natl Acad Sci U S A 113:12150–12155.

8. Trinquier A, Ulmer JE, Gilet L, Figaro S, Hammann P, Kuhn L, Braun F, Condon C. 2019. tRNA Maturation Defects Lead to Inhibition of rRNA Processing via Synthesis of pppGpp. Mol Cell 74:1227–1238 e3.

9. Ishimura R, Nagy G, Dotu I, Zhou H, Yang XL, Schimmel P, Senju S, Nishimura Y, Chuang JH, Ackerman SL. 2014. RNA function. Ribosome stalling induced by mutation of a CNS-specific tRNA causes neurodegeneration. Science 345:455–9.

10. Nicolas P, Mader U, Dervyn E, Rochat T, Leduc A, Pigeonneau N, Bidnenko E, Marchadier E, Hoebeke M, Aymerich S, Becher D, Bisicchia P, Botella E, Delumeau O, Doherty G, Denham EL, Fogg MJ, Fromion V, Goelzer A, Hansen A, Hartig E, Harwood CR, Homuth G, Jarmer H, Jules M, Klipp E, Le Chat L, Lecointe F, Lewis P, Liebermeister W, March A, Mars RA, Nannapaneni P, Noone D, Pohl S, Rinn B, Rugheimer F, Sappa PK, Samson F, Schaffer M, Schwikowski B, Steil L, Stulke J, Wiegert T, Devine KM, Wilkinson AJ, van Dijl JM, Hecker M, Volker U, Bessieres P, et al. 2012. Condition-dependent transcriptome reveals high-level regulatory architecture in Bacillus subtilis. Science 335:1103–6.

11. DeLoughery A, Lalanne JB, Losick R, Li GW. 2018. Maturation of polycistronic mRNAs by the endoribonuclease RNase Y and its associated Y-complex in Bacillus subtilis. Proc Natl Acad Sci U S A 115:E5585–E5594.

12. Kery MB, Feldman M, Livny J, Tjaden B. 2014. TargetRNA2: identifying targets of small regulatory RNAs in bacteria. Nucleic Acids Res 42:W124–9.

13. Murray HD, Schneider DA, Gourse RL. 2003. Control of rRNA expression by small molecules is dynamic and nonredundant. Mol Cell 12:125–34.

14. VanBogelen RA, Kelley PM, Neidhardt FC. 1987. Differential induction of heat shock, SOS, and oxidation stress regulons and accumulation of nucleotides in Escherichia coli. J Bacteriol 169:26–32.

15. Kriel A, Bittner AN, Kim SH, Liu K, Tehranchi AK, Zou WY, Rendon S, Chen R, Tu BP, Wang JD. 2012. Direct regulation of GTP homeostasis by (p)ppGpp: a critical component of viability and stress resistance. Mol Cell 48:231–41.

16. Durand S, Gilet L, Condon C. 2012. The essential function of B. subtilis RNase III is to silence foreign toxin genes. PLoS Genet 8:e1003181.

17. Deana A, Belasco JG. 2005. Lost in translation: the influence of ribosomes on bacterial mRNA decay. Genes Dev 19:2526–33.

18. Rhaese HJ, Dichtelmuller H, Grade R. 1975. Studies on the control of development. Accumulation of guanosine tetraphosphate and pentaphosphate in response to inhibition of protein synthesis in Bacillus subtilis. Eur J Biochem 56:385–92.

19. Cashel M. 1969. The control of ribonucleic acid synthesis in Escherichia coli. IV. Relevance of unusual phosphorylated compounds from amino acid-starved stringent strains. J Biol Chem 244:3133–41.

20. Kurland CG, Maaloe O. 1962. Regulation of ribosomal and transfer RNA synthesis. J Mol Biol 4:193–210.

21. Katz A, Elgamal S, Rajkovic A, Ibba M. 2016. Non-canonical roles of tRNAs and tRNA mimics in bacterial cell biology. Mol Microbiol 101:545–58.

22. Raina M, Ibba M. 2014. tRNAs as regulators of biological processes. Front Genet 5:171.

23. Lee YS, Shibata Y, Malhotra A, Dutta A. 2009. A novel class of small RNAs: tRNA-derived RNA fragments (tRFs). Genes Dev 23:2639–49.

24. Kumar P, Kuscu C, Dutta A. 2016. Biogenesis and Function of Transfer RNA-Related Fragments (tRFs). Trends Biochem Sci 41:679–689.

25. Kim HK, Fuchs G, Wang S, Wei W, Zhang Y, Park H, Roy-Chaudhuri B, Li P, Xu J, Chu K, Zhang F, Chua MS, So S, Zhang QC, Sarnow P, Kay MA. 2017. A transfer-RNA-derived small RNA regulates ribosome biogenesis. Nature 552:57–62.

26. Mohanty BK, Kushner SR. 2022. Inactivation of RNase P in Escherichia coli significantly changes post-transcriptional RNA metabolism. Mol Microbiol 117:121–142.

27. Bechhofer DH, Oussenko IA, Deikus G, Yao S, Mathy N, Condon C. 2008. Analysis of mRNA decay in Bacillus subtilis. Methods Enzymol 447:259–76.

28. Wegscheid B, Condon C, Hartmann RK. 2006. Type A and B RNase P RNAs are interchangeable in vivo despite substantial biophysical differences. EMBO Rep 7:411–7.

29. Koo BM, Kritikos G, Farelli JD, Todor H, Tong K, Kimsey H, Wapinski I, Galardini M, Cabal A, Peters JM, Hachmann AB, Rudner DZ, Allen KN, Typas A, Gross CA. 2017. Construction and Analysis of Two Genome-Scale Deletion Libraries for Bacillus subtilis. Cell Syst 4:291–305 e7.

30. Chary VK, Amaya EI, Piggot PJ. 1997. Neomycin- and spectinomycin-resistance replacement vectors for Bacillus subtilis. FEMS Microbiol Lett 153:135–9.

31. Gendron N, Putzer H, Grunberg-Manago M. 1994. Expression of both *Bacillus subtilis* threonyl-tRNA synthetase genes is autogenously regulated. J Bacteriol 176:486–494.

