## Supplementary Tables and Figures for "Effect of tRNA maturase depletion on the levels and stabilities of ribosome assembly cofactor mRNAs in *Bacillus subtilis*"

**Table S1: *B. subtilis* strains used in this study**

| Strains | Genotype | Reference |
|---|---|---|
| SSB318 | W168 *Pspac-rnpB:pMUTIN ery* | (28) |
| SSB1002 | W168 *trp+* | Lab strain |
| CCB288 | W168 *amyE::Pspac-rnc Cm rnc::spc pMAP65* | (16) |
| CCB321 | W168 *Pspac-rnz:pMUTIN ery* pMAP65 *kan* | (8) |
| CCB418 | W168 *txpA -10Δ yonT::ery rnc::spc* | (16) |
| CCB504 | W168 *Pxyl-rnpA Cm* | (8) |
| CCB664 | W168 *rimM*::*ery* | (29); This study |
| CCB994 | W168 *Pspac-rnpB:pMUTIN ery amyE::pHM2-ylqD-rimM Cm* | This study |
| CCB1008 | W168 *Pspac-rnpB:pMUTIN ery amyE::pHM2-ylqD*-rimM Cm* | This study |
| CCB1014 | W168 *Pspac-rnpB:pMUTIN ery amyE::pHM2 Cm* | This study |
| CCB1017 | W168 *Pspac-rnpB:pMUTIN ery amyE::pHM2-rimM Cm* | This study |
| CCB1031 | W168 *Pxyl-rnpA Cm amyE::pHM2-rimM Spc* | This study |
| CCB1050 | W168 *yjbM::spc ywaC::kan relA::ery* | (8) |
| CCB1057 | W168 *yjbM::spc ywaC::kan Pxyl-rnpA Cm relA::ery* | (8) |
| CCB1125 | W168 *yjbM::spc ywaC::kan amyE::pX-ywaC Cm relA::ery* | (8) |
| CCB1137 | W168 *yjbM::spc ywaC::kan Pspac-rnpB:pMUTIN relA::ery::tet* | (8) |

**Table S2. Details of strain construction**

| Strain number | Plasmid | Description | Source/Ref. |
|---|---|---|---|
| CCB994 | 752 | Plasmid 752 linearized with XhoI and integrated into SSB318 *amyE* locus. | This study; (8) |
| CCB1008 | 762 | Plasmid 762 linearized with XhoI and integrated into SSB318 *amyE* locus. | This study; (8) |
| CCB1014 | 606 | Plasmid 606 linearized with XhoI and integrated into SSB318 *amyE* locus. | This study; (8) |
| CCB1017 | 764 | Plasmid 764 linearized with SpeI and integrated into SSB318 *amyE* locus. | This study; (8) |
| CCB 1031 | Plasmid 764 and pCS | First, plasmid 764 linearized with SpeI was integrated into *amyE* in WT. Then the plasmid 764 Cm cassette was switched to Spc using an antibiotic cassette switching vector (pCS). The resulting strain was then transformed with CCB504 gDNA. | This study; (8, 30) |

**Table S3: Plasmids constructed for this study**

| Plasmid number | Plasmid name | Initial vector | Insert | Oligos for PCR | Description | Source /Ref. |
|---|---|---|---|---|---|---|
| 606 | pHM2-Pspac(con) | pHM2 | Pspac(con) | | Promoter $P_{spac}$ from pMUTIN-4M cloned without the operator sequence using EcoRI-HindIII restriction sites | This study; (31) |
| 752 | pHM2-Pspac(con) -ylqD-rimM | 606 | *ylqD-rimM* | CC1985 + CC1986 | Insert amplified from gDNA and cloned in BamHI/SalI under plasmid 606 constitutive promoter. | This study |
| 762 | pHM2-Pspac(con) -ylqD*-rimM | 606 | *ylqD*-rimM* | CC1985 + CC2012; CC2011 + CC1986 | ylqD* and *rimM* were amplified from plasmid 752. The CC2011/CC2012 oligos introduce mutations in *ylqD* ORF. *ylqD*-rimM* overlap was obtained by overlap PCR with underlined oligos and cloned in BamHI/SalI. | This study |
| 764 | pHM2-Pspac(con) -rimM | 606 | *rimM* | CC1986 + CC2034 | Insert amplified from gDNA and cloned in BamHI/XhoI under plasmid 606 constitutive promoter. | This study |

**Table S4: Oligonucleotides used in this study**

| Oligo | Gene | Sequence |
|---|---|---|
| CC1005 | *rnpA* | ACTGGCGGTTTGCAACTGATGTCCC |
| CC1006 | *rnpB* | TGCGAGCATGGACTTTCCTCTACAG |
| CC1560 | *rpsU* | CTAGCAGCTTCAGACTTTTTCTTGCGC |
| CC1841 | *ylxS* | GAGTTCCAGCTGAAGGCTGTCTACGATTGGC |
| CC1842 | *rbfA* | GCTTTTGCCAGCCCTTTCAGCGCTTCTTCCC |
| CC1845 | *rimM* | GAAATCACCCGCACTTCGCCTTTGATTCCGTG |
| CC1846 | *era* | CCTTGTTTCTCGTCGTTTGGGGCTTATCGC |
| CC1847 | *yqeH* | CGACCAGAGAGTCCGTTTCTCCAATACCGTG |
| CC1915 | trnJ-lys | GACTCGAACCTTCGACCCTCTGATTAAAAG |
| CC1985 | ylqD fw | TAAGGATCCGCTTAGGCAAAATAGAAAAGCCTGGACGAG |
| CC1986 | rimM rvs | TAACTCGAGAAAAAGGCCATCCGTCAGGATGGCCGAGCACGCCTTCAAACATTTaGGGAAACAGCG |
| CC2011 | ylqD* fw | GGGGATGACTGGCATcAGtttcaccCAGCGAACACCATCGTC |
| CC2012 | ylqD* rvs | GACGATGGTGTTCGCTGggtgaaaCTgATGCCAGTCATCCCC |
| CC2034 | rimM fw | ATATGGATCCTGGAAGAGGTGATCATATGACAAAGCGATGG |
| CC2099 | *ylqD* | CTCGGTTAAGACTTGCATAACGGCTACACGG |
| CC2100 | *ylqC* | GTCATCTGGATGATCAACAAGCGGCGTCAC |
| CC2101 | *trmD* | CTGCCTTTGATGTCAGGTCCTCGACCGCGTC |
| CC2103 | *ffh* | GAAATCGTCTGCTGCAGTCGGTCGGCTAATC |
| CC2143 | *rpsP* | GATGAAACGGCCGTCACGTGGTGAACGAGAATC |
| CC2144 | *rplS* | CAGGACGGAACGCAGGAAGATCAGTACGAAG |
| CC2197 | *yjcK* | CGAAACCGTCCCGATTAATCTATCGTCTGACG |
| CC2198 | *ydaF* | GATTGTTTCTCGGTACGTGTCAGCGCTGCTTG |
| CC2199 | *yfmL* | GACGAGCTGATCCGTTTCATCAAGCACGATC |
| CC2200 | *cpgA fwd* | CAACGAGCTTATCAGGCCGCCAATTTGCAAC |
| CC2201 | *cpgA + PT7 rvs* | gctctaatacgactcactatagggACGTGTGAATCAGCTCCACGTGGCGGG |
| CC2213 | *ywaA* | GATGCAGAGGCGGTCGTTTGATTGATTCAG |
| CC2215 | *ilvA* | CACATCCGGATCATCGAACGGATGGATAAACGTC |
| CC2450 | *ksgA* | CTGTTTTCTCCGTCACTTCCGCGTGATCAAC |

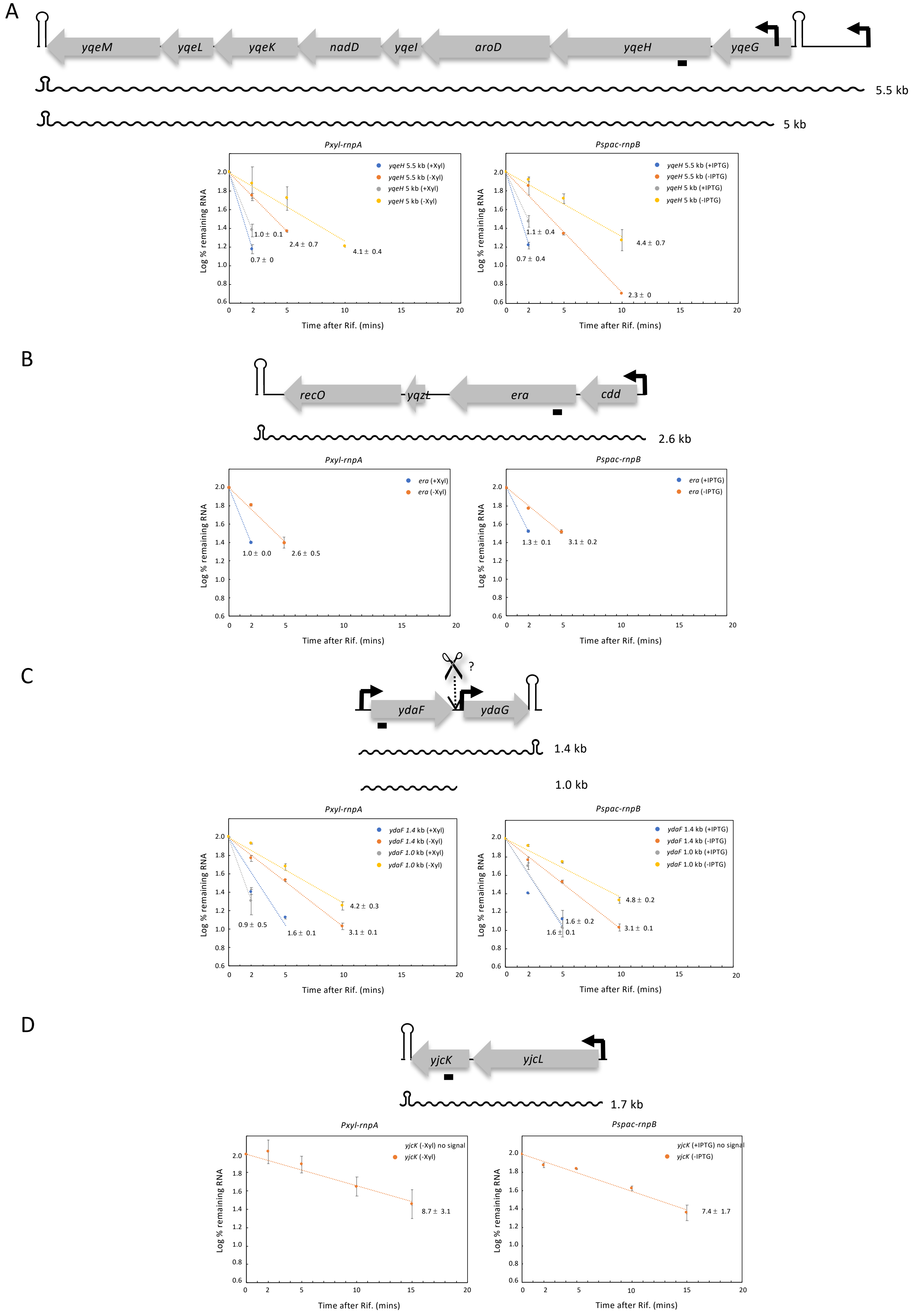


Figure S1: Schematics and RNA decay plots of up-regulated transcripts in RnpA and RnpB-depletion strains. Open reading frames (ORFs; not to scale) are shown as gray arrows and transcripts as wavy lines. Sizes are as indicated. Promoters are represented by black arrows and terminators as hairpins. Suspected endoribonucleolytic cleavage sites are indicated by a scissors symbol with a question mark. Quantifications are from the average of two experiments, with half-lives as indicated.

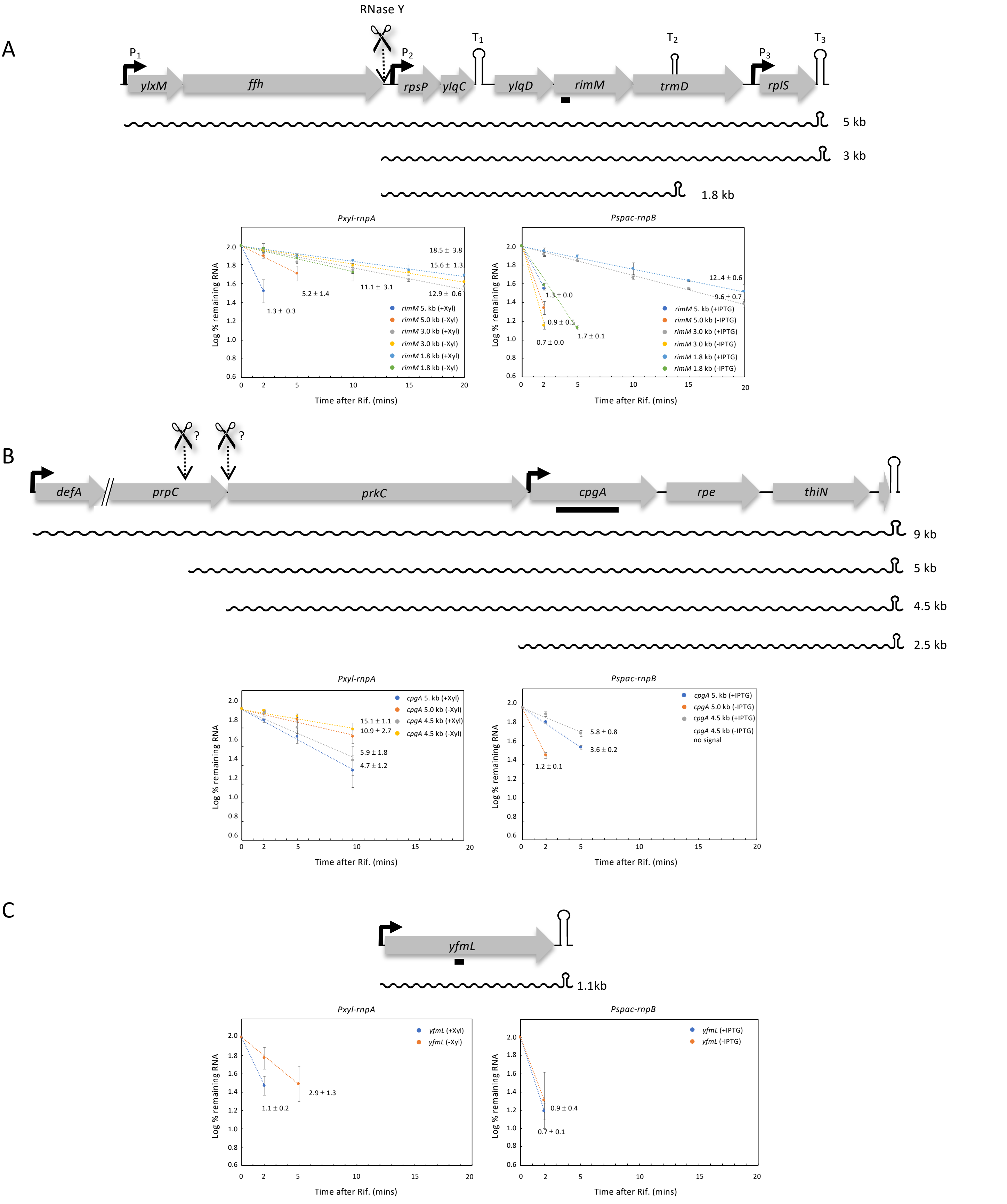


Figure S2: Schematics and RNA decay plots of down-regulated transcripts in RnpA and RnpB-depletion strains. Open reading frames (ORFs; not to scale) are shown as gray arrows and transcripts as wavy lines. Sizes are as indicated. Promoters are represented by black arrows and terminators as hairpins. Suspected endoribonucleolytic cleavage sites are indicated by a scissors symbols with question marks. Quantifications are from the average of two experiments, with half-lives as indicated.

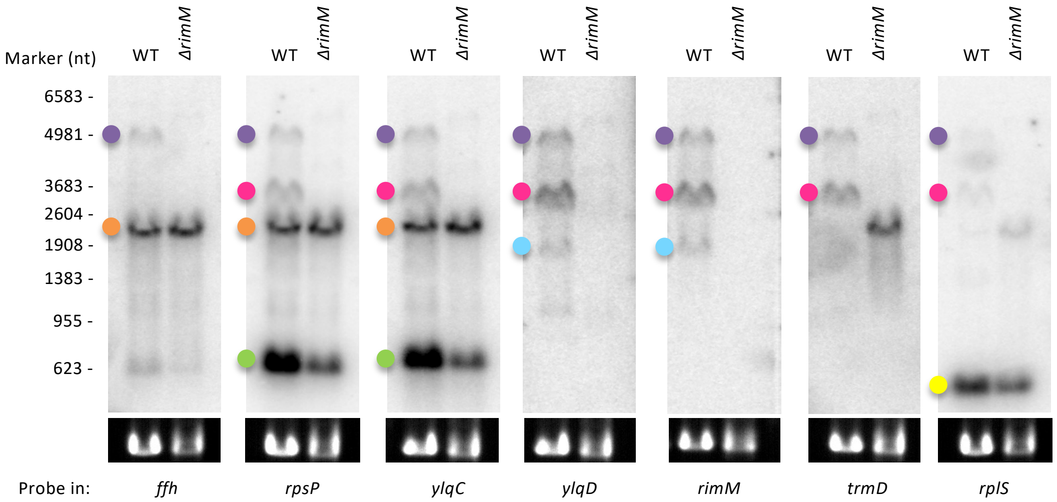


Figure S3: Northern blot analysis to determine the structure of the *rimM* operon. Total RNA from WT or *ΔrimM* strains was probed with oligonucleotides targeting different ORFs of the *rimM* operon (indicated below each panel). 16S rRNA levels (ethidium bromide stained) are shown as a loading control. The *ffh* and *ylqC* blots were stripped and reprobed for *rplS* and *ylqD*, respectively. Colored dots correspond to the color of transcripts shown in Figure 4A.

| | levels | | | stability | | Cm | | ppGpp° | | ppGpp⁺ | growth rate |
|---|---|---|---|---|---|---|---|---|---|---|---|
| | *rnz* | *rnpA* | *rnpB* | *rnpA* | *rnpB* | sub | MIC | *rnpA* | *rnpB* | - | *rnc* |
| FL | - | - | ↓ | ↑ | - | ↷ | ↷ | ↑ | ↑ | ↓ | ↓ |
| Y/$P_2$-$T_3$ | ↓ | ↓ | ↓ | - | ↓ | ↷ | ↷ | - | ↑ | ↓ | ↓ |
| Y/$P_2$-$T_2$ | ↓ | ↓ | - | - | ↓ | ↷ | ↷ | ↑ | ↑ | - | ↓ |

Figure S4. Summary of effects on 3 three *rimM* containing transcripts. Data are taken from Northern blots shown in Figures 1 (levels), 2-3 (stability, Cm), 6A (ppGpp$^0$), 6B (ppGpp$^+$) and 7 (growth rate). Depleted tRNA maturases are indicated. Transcripts are listed according to size: full-length (FL), RNase Y/$P_2$-$T_3$ and RNase Y/$P_2$-$T_2$. Red arrows show down-effects, green arrows show up-effects, (-) no effect. Curved arrow in sub-inhibitory Cm condition (sub) reflects initial increase followed by decrease.

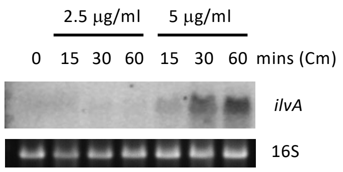


Figure S5. Induction of expression of the CodY regulon member *ilvA* by chloramphenicol. Northern blots of total RNA isolated at different times after addition of 0.5x MIC and MIC of chloramphenicol (Cm). 16S rRNA levels (ethidium bromide stained) are shown as a loading control.
